# Collaborative binding by diverse pioneer transcription factors amplifies Polycomb repression to preserve lineage fidelity

**DOI:** 10.64898/2026.05.19.726322

**Authors:** Gerardo Mirizio, Morgan Buckley, Katie Ludwig, Satoshi Matsui, Samuel Sampson, Hee-Woong Lim, Makiko Iwafuchi

**Affiliations:** Division of Developmental Biology, Center for Stem Cell & Organoid Medicine (CuSTOM), Cincinnati Children’s Hospital Medical Center, Department of Pediatrics, University of Cincinnati College of Medicine, 3333 Burnet Ave., Cincinnati, OH 45229, USA; Division of Biomedical Informatics, Cincinnati Children’s Hospital Medical Center, Department of Pediatrics, University of Cincinnati College of Medicine, 3333 Burnet Ave., Cincinnati, OH 45229, USA

## Abstract

Cooperative and collaborative modes of transcription factor (TF) binding are central to eukaryotic gene activation, providing regulatory specificity and robustness in cell fate control. However, gene activation is only half the story of transcriptional regulation, and cells also require mechanisms to repress inappropriate gene programs. It remains largely unknown how broadly expressed repressive TFs and epigenetic regulators achieve context-specific repression required for proper cell differentiation. Here, we show that the repressive TF PRDM1 cooperates with lineage- and stage-specific pioneer TFs during human endoderm differentiation. Notably, the number of diverse pioneer TFs bound at enhancers determines the strength of PRDM1-mediated enhancer repression. Mechanistically, collaborative binding by multiple pioneer TFs synergistically promotes local nucleosome remodeling and stabilizes PRDM1 occupancy, culminating in the formation of “hyper-bivalent” enhancers that are characterized by strongly elevated levels of H3K4me1 and Polycomb-associated modifications. These enhancers reinforce the repression of alternative-lineage, and past and future developmental programs. Together, our findings redefine the roles of TF cooperation and chromatin accessibility beyond the prevailing activation-centric view. We propose a “collaborative repression” model in which diverse pioneer TFs establish focal chromatin accessibility that reinforces the context-specific assembly of repressive regulators, thereby safeguarding lineage and developmental fidelity.

**Highlights:**

- PRDM1 cooperates with diverse lineage- and stage-specific pioneer TFs
- The number of pioneer TFs bound at enhancers determines the strength of Polycomb repression
- Collaborative binding by diverse pioneer TFs and PRDM1 establishes hyper-bivalent enhancers
- Collaborative repression safeguards lineage and developmental fidelity

## INTRODUCTION

Both cooperative and collaborative modes of binding among multiple transcription factors (TFs), mediated by direct TF–TF interactions or indirect mechanisms, drive context-specific enhancer activation.^1–4^ Functionally, multiple TF-binding sites within an enhancer promote stable TF occupancy and increase regulatory specificity, enabling robust lineage-specific gene expression.^5–13^ Central to establishing these regulatory platforms are lineage-specific pioneer TFs. They engage nucleosomal DNA within compacted chromatin and locally open target sites.^14–17^ This pioneering activity facilitates the recruitment of additional TFs and epigenetic regulators, including H3K4 methyltransferases, which collectively promote gene activation.^18–22^

Equally important, lineage-specific gene regulation requires repressive mechanisms to prevent inappropriate activation of off-target genes. During development, the Polycomb repressive complexes (PRCs) play a critical role in repressing alternative-lineage programs.^23–25^ PRC1 and PRC2 deposit the Polycomb-associated histone modifications, H2AK119ub1 and H3K27me3, respectively, at enhancers and CpG-rich promoters of key developmental genes. These modifications frequently coexist with H3K4 methylation, forming bivalent chromatin states that are transcriptionally repressed yet poised for activation.^26–29^ While PRC recruitment to promoters has been attributed to RNA-mediated pathways and hypomethylated CpG islands, how PRCs are selectively recruited to enhancers remains less well understood.^30,31^ Several studies indicate that the repressive TF REST, which is broadly expressed across non-neural tissues, facilitates PRC recruitment.^32,33^ We and others have also shown that the PR domain zinc finger TF PRDM1 contributes to Polycomb-associated epigenetic repression in foregut endoderm, primordial germ cells, and plasma cells, with PRDM1 regulating distinct gene sets in each cellular context.^34–36^ Therefore, a key question remains: given that repressive TFs and PRCs are broadly expressed, how do they achieve context-specific repression?

To address this gap, we used human endoderm differentiation as a tractable model, during which PRDM1 is transiently upregulated in definitive endoderm and foregut endoderm and downregulated thereafter. We first find that PRDM1 dynamically shifts its genomic occupancy from definitive endoderm to foregut endoderm. While we previously showed that PRDM1 cooperates with FOXA pioneer TFs in foregut endoderm,^35^ FOXA TFs are not yet highly expressed at the definitive endoderm stage. Instead, PRDM1 co-localizes with the diverse pioneer TFs EOMES, GATA6, and SOX17 in definitive endoderm, predominantly at Polycomb-associated enhancers. Using a doxycycline-inducible CRISPRi system, we show that *PRDM1* knockdown in definitive endoderm results in de-repression of genes associated with alternative-lineage programs, accompanied by reduced enrichment of Polycomb-associated histone modifications. A lentivirus-based massively parallel reporter assay (lentiMPRA) reveals that the number of diverse pioneer TF-binding sites within regulatory elements determines the strength of PRDM1-mediated repression. Mechanistically, chromatin accessibility at PRDM1-bound sites and enrichment of bivalent histone modifications in flanking regions progressively increase with the number and diversity of co-bound pioneer TFs, culminating in the formation of “hyper-bivalent” enhancers that are characterized by strongly elevated levels of H3K4me1 and Polycomb-associated modifications. These hyper-bivalent enhancers reinforce the repression of genes specific not only to alternative lineages, but also to past and future endoderm developmental stages.

Together, these findings redefine our understanding of TF cooperation and chromatin accessibility, extending them beyond the prevailing activation-centric view. We propose a “collaborative repression” model, in which collaborative binding by diverse lineage- and stage-specific pioneer TFs establishes focal chromatin accessibility that reinforces the context-specific recruitment of repressive regulators. This collaborative repression mechanism prevents alternative-lineage differentiation, restrains reactivation of earlier developmental programs, and limits precocious activation of later programs, thereby safeguarding lineage and developmental fidelity.

## RESULTS

### PRDM1 cooperates with stage- and lineage-specific pioneer TFs to establish Polycomb-associated enhancers

To determine how PRDM1 achieves context-specific repression, we used a human pluripotent stem cell (hPSC) differentiation model that generates homogeneous populations of anterior primitive streak (APS), definitive endoderm (DE), posterior foregut (PFG), and liver bud progenitor (LBP) cells (**Figure 1A** and **Table S1**).^35,37^ RNA sequencing (RNA-seq) analysis showed that PRDM1 was induced upon exit from pluripotency, peaked at the DE and PFG stages, and then declined during differentiation into LBP cells (**Figure 1B**). To assess PRDM1 genomic occupancy in DE and PFG cells, where it is highly expressed, we performed chromatin immunoprecipitation followed by sequencing (ChIP-seq) for PRDM1. Although these stages are consecutive, PRDM1 dynamically shifted its genomic occupancy from DE to PFG (**Figure 1C**). Motif analysis of stage-specific PRDM1-bound sites revealed distinct motif enrichment. PRDM1-bound sites in DE cells were enriched for EOMES, GATA, ZIC, and SOX17 motifs (**Figure 1D**, DE-specific). EOMES (T-box family), GATA6 (GATA family), ZIC2 (Zinc finger of the cerebellum family), and SOX17 (SRY-related HMG-box family) are expressed in DE cells and have all been reported to function as pioneer TFs.^9,13,18,20,38–41^ In contrast, PRDM1-bound sites in PFG cells were most strongly enriched for a FOXA pioneer TF motif, consistent with our previous finding,^35^ followed by TEAD, GATA, and SOX4 motifs (**Figure 1D**, PFG-specific). These results suggest that PRDM1 cooperates with stage-specific combinations of pioneer TFs to achieve its context-specific genomic occupancy.

**Figure 1.**
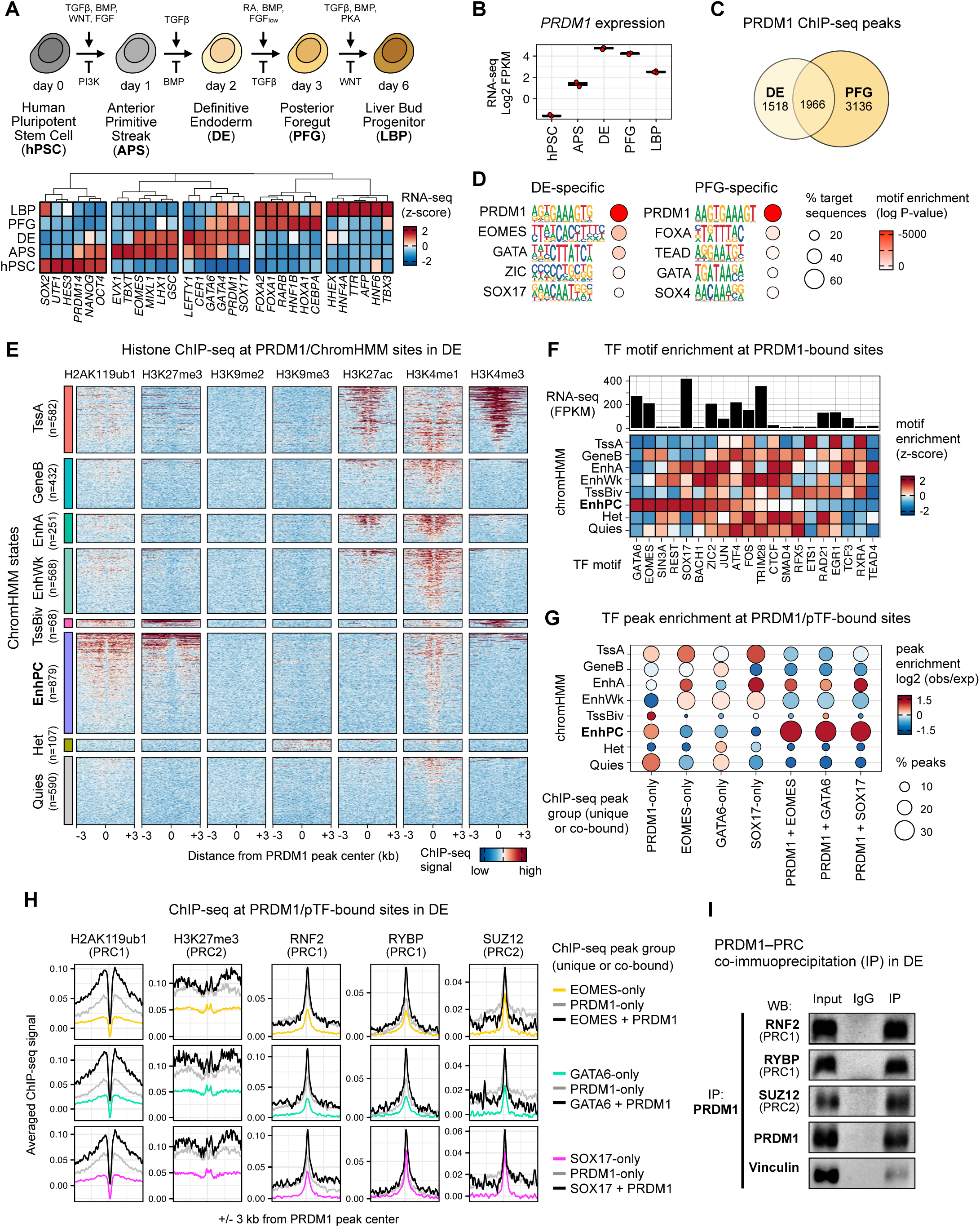
PRDM1 cooperates with stage- and lineage-specific pioneer TFs to establish Polycomb-associated enhancers. **(A)** Schematic representation of the differentiation protocol from human pluripotent stem cells (hPSCs) to endodermal cells, and expression matrix of marker genes derived from RNA-seq data (*n* = 3 biological replicates) across differentiation stages. **(B)** PRDM1 expression dynamics during endoderm differentiation (RNA-seq, *n* = 3 biological replicates). **(C)** Intersection of PRDM1 ChIP-seq peaks in DE and PFG (*n* = 2 biological replicates). **(D)** Top motifs enriched at DE-specific, DE-PFG common, and PFG-specific PRDM1 peaks by HOMER. *P-*values were calculated using a cumulative hypergeometric distribution test. **(E)** Heatmap of histone ChIP-seq signal centered on PRDM1 peaks and clustered by ChromHMM categories at the DE stage. TssA: active promoters, GeneB: gene bodies, EnhA: active enhancers, EnhWk: weak enhancers, TssBiv: bivalent promoters, EnhPC: Polycomb-associated enhancers, Het: heterochromatin, Quies: quiescent regions. **(F)** Heatmap of transcription factor (TF) motif enrichment at PRDM1 peaks across ChromHMM states, rank-ordered by TF motif enrichment at PRDM1/EnhPC sites. Z-scores indicate relative motif enrichment across chromatin states. **(G)** Dot plot showing enrichment of ChIP-seq peak groups across ChromHMM states. The size of the dots represents the proportion of peaks enriched in each ChromHMM state. Log2 (obs/exp) stands for the base-2 logarithm of the observed-to-expected ratio. **(H)** Averaged plots of ChIP-seq signal in DE cells for PRC1 (H2AK119ub1) and PRC2 (H3K27me3) histone modifications as well as PRC1 (RING1B, RYBP) and PRC2 (SUZ12) subunits, across sites bound by PRDM1 alone, pioneer TFs alone, or co-bound by PRDM1 and pioneer TFs (*n* = 2 biological replicates). **(I)** Immunoprecipitation (IP) of PRDM1 followed by Western blot (WB) for RNF2, RYBP, and SUZ12 in nuclear extracts from DE cells (*n* = 3 biological replicates). WB for PRDM1 and Vinculin was performed as positive and negative controls, respectively.

To comprehensively characterize PRDM1-bound regions, we focused on the DE stage, when PRDM1 is highly induced, and annotated PRDM1 peaks into distinct chromatin states. We performed ChIP-seq for epigenetic modifications associated with active regulatory elements (H3K27ac), active or primed enhancers (H3K4me1), active or primed promoters (H3K4me3), PRC2 (H3K27me3), PRC1 (H2AK119ub1), and heterochromatin (H3K9me2/3) in DE cells. We then classified PRDM1 peaks using Chromatin Hidden Markov Model (ChromHMM) annotations generated in hPSC-derived endoderm cells by the ENCODE Project (**Figure 1E**).^42^ This enabled us to interpret PRDM1-bound sites and our epigenomic profiles within a unified classification of functional chromatin states. PRDM1-bound sites were annotated into eight distinct ChromHMM states corresponding to active promoters (TssA; n = 582), gene bodies (GeneB; n = 432), active enhancers (EnhA; n = 251), weak enhancers (EnhWk; n = 568), bivalent promoters (TssBiv; n = 68), Polycomb-associated enhancers (EnhPC; n = 879), heterochromatin (Het; n = 107), and quiescent regions (Quies; n = 590) (**Figure 1E**). Notably, a substantial fraction of PRDM1-bound sites localized to Polycomb-associated enhancers, characterized by enrichment of the PRC-associated modifications H2AK119ub1 and H3K27me3, together with weak enrichment of H3K4me1 (**Figure 1E**, EnhPC).

Having classified PRDM1-bound sites by chromatin state, we next sought to identify TF candidates that cooperate with PRDM1 within each chromatin state. We thus performed differential motif enrichment analysis of PRDM1-bound sites stratified by ChromHMM state and considered the expression levels of the corresponding TFs in DE cells (**Figure 1F**). PRDM1-bound active and weak enhancers were enriched for TF motifs recognized by ZIC2, as well as motifs for downstream effectors of FGF and TGF-β signaling pathways, such as FOS/JUN (AP-1 complex) and SMAD4. In contrast, PRDM1-bound Polycomb-associated enhancers were enriched for motifs recognized by the pioneer TFs, GATA6, EOMES, and SOX17 (**Figure 1F**). Given their robust expression at the DE stage (**Figure 1F**), we examined their co-occupancy with PRDM1 at Polycomb-associated enhancers by performing ChIP-seq for these pioneer TFs in DE cells. We classified EOMES, GATA6, and SOX17 peaks as either pioneer TF-only sites or PRDM1+pioneer TF co-bound sites based on their overlap with PRDM1 peaks within a 200-bp window (**Figure S1A**). Pioneer TF-only sites were preferentially enriched at weak/active enhancers and active promoters, whereas PRDM1+pioneer TF co-bound sites were preferentially enriched at Polycomb-associated enhancers (**Figures 1G** and **S1B**). These results suggest that PRDM1 cooperates with EOMES, GATA6, and SOX17 to target DE-specific, Polycomb-associated enhancers.

To investigate whether PRDM1 and the DE-enriched pioneer TFs cooperate to recruit PRCs and establish Polycomb-associated chromatin states, we performed ChIP-seq for the PRC1 catalytic subunit RNF2 (RING1B), the non-canonical PRC1 subunit RYBP, the PRC2 core subunit SUZ12, and the PRC-associated histone modifications H2AK119ub1 and H3K27me3. We then assessed their enrichment at pioneer TF-only, PRDM1-only, and PRDM1+pioneer TF co-bound sites (**Figure 1H**). PRC subunits and PRC-associated modifications showed minimal enrichment at pioneer TF-only-bound sites, greater enrichment at PRDM1-only-bound sites, and the strongest enrichment at PRDM1+pioneer TF co-bound sites (**Figure 1H**). Importantly, PRC-associated modifications showed little or no enrichment at regions co-occupied by different pioneer TFs in the absence of PRDM1, indicating a critical role for PRDM1 in efficient PRC recruitment (**Figure S1C**). Enrichment of PRC1 subunits, but not the PRC2 subunit SUZ12, was further increased at PRDM1+pioneer TF sites relative to PRDM1-only sites, suggesting that pioneer TFs reinforce PRDM1-mediated recruitment or stabilization of PRC1 (**Figure 1H**). To test whether PRDM1 physically interacts with PRCs in DE cells, we performed co-immunoprecipitation (co-IP) using endogenous nuclear extracts treated with Benzonase nuclease to minimize interactions arising from DNA-mediated proximity. Indeed, PRDM1 co-immunoprecipitated with the PRC1 subunits RNF2 and RYBP and the PRC2 subunit SUZ12 (**Figure 1I**). Altogether, these results indicate that PRDM1 cooperates with diverse stage- and lineage-specific pioneer TFs to recruit and/or stabilize PRCs and establish Polycomb-associated enhancers during endoderm differentiation.

### PRDM1 safeguards endoderm identity by Polycomb-mediated repression of alternative-lineage genes

To define the gene regulatory function of PRDM1 during early endoderm differentiation, we knocked down *PRDM1* (*PRDM1*-KD) in DE cells with a doxycycline (Dox)-inducible CRISPR interference (CRISPRi) system (**Figure 2A**). Dox treatment for 2 days, starting at the onset of differentiation, efficiently reduced PRDM1 protein levels in DE cells (**Figures 2B** and **S2A**). RNA-seq analysis of control and *PRDM1*-KD cells identified 570 differentially expressed genes (FC ≥ 1.5; FDR ≤ 0.05) (**Figure 2C**, **Table S2**). Gene Ontology (GO) analysis of up-regulated genes upon *PRDM1* knockdown revealed enrichment of mesoderm-associated developmental programs (e.g., muscle cell fate, heart morphogenesis, cartilage development, and ossification) and ectoderm-associated programs (e.g., synapse assembly, synaptic transmission, and cell migration) (**Figure S2B**). To further characterize changes in cell identity in *PRDM1*-KD cells, we compared their transcriptional profile with those of other major cell types at comparable early human developmental stages using CellNet,^43^ a gene regulatory network (GRN)-based computational framework that quantitatively assesses cell identity. To this end, we built a reference atlas of cell types spanning early hPSC differentiation from more than 500 RNA-seq datasets and used this atlas to train lineage-specific GRN classifiers (**Figure S1D** and **S1E**). CellNet classification of query samples against the reference cell types revealed that *PRDM1*-KD cells exhibited lower DE identity scores and higher scores for mesendoderm (corresponding to anterior primitive streak [APS] and mid-primitive streak [MPS]), mesoderm (paraxial mesoderm [PM] and lateral plate mesoderm [LPM]), and neuroectoderm (NE) cell types as compared to control cells (**Figure 2D**). These results suggest that PRDM1 safeguards endoderm identity by restraining alternative lineage-specific transcriptional networks and preventing reactivation of earlier developmental programs.

**Figure 2.**
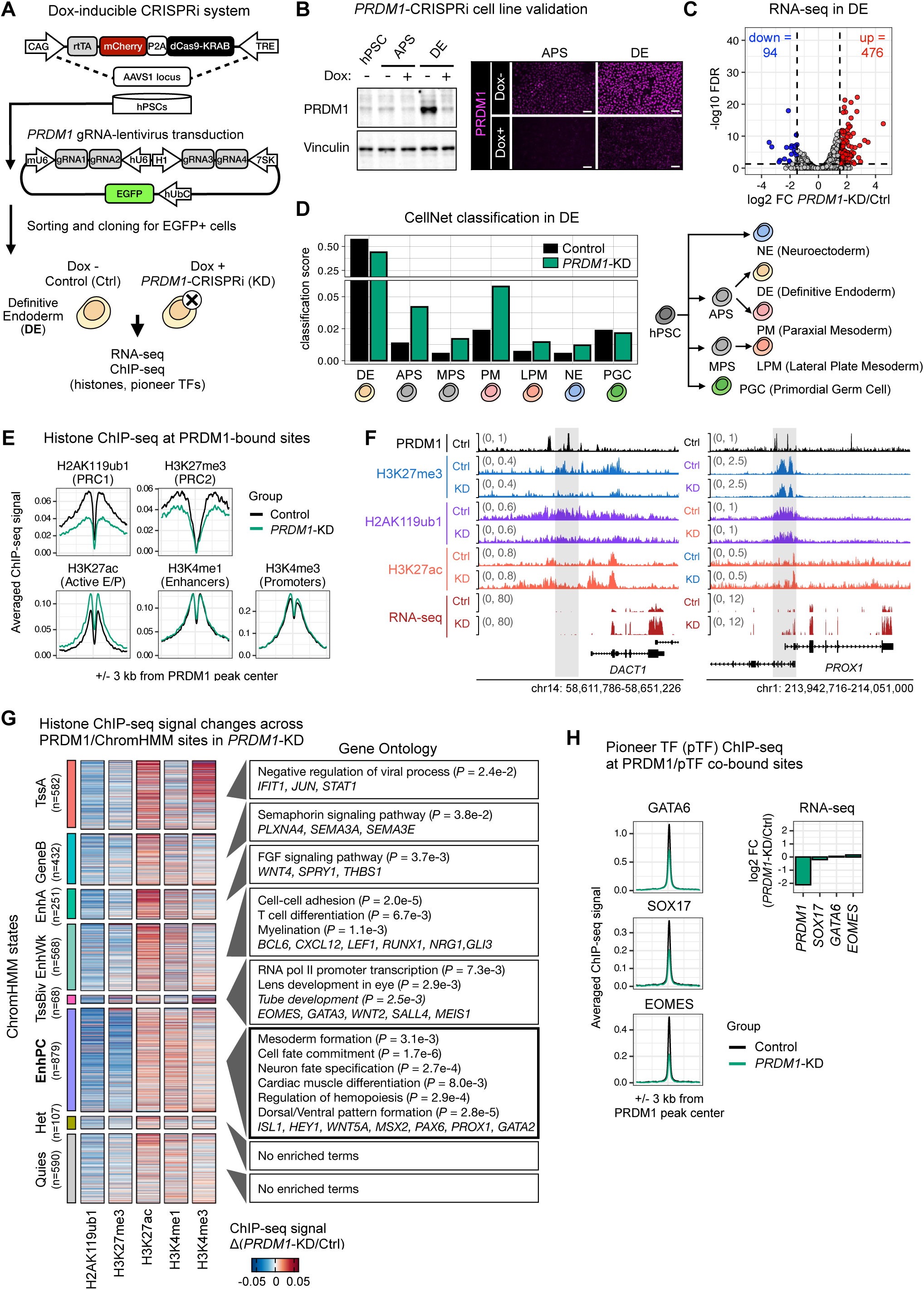
PRDM1 safeguards endoderm identity by Polycomb-mediated repression of alternative-lineage genes. **(A)** Schematic representation of the Dox-inducible CRISPRi system used to knock down *PRDM1* expression. Guide RNAs targeting *PRDM1* are constitutively expressed, whereas dCas9 is induced upon doxycycline (Dox) administration. **(B)** Assessment of PRDM1 knockdown efficiency in *PRDM1*-CRISPRi cells by Western blot and immunofluorescence during differentiation from hPSC to DE cells. Scale bar = 100 µm. **(C)** Volcano plot of RNA-seq data from control and *PRDM1*-KD cells at the DE stage (*n* = 2 biological replicates). Differential expression was assessed using the Wald test, with *P-*values corrected for multiple testing using the Benjamini–Hochberg method (FDR ≤ 0.05) and an absolute log2 fold change ≥ 1.5. **(D)** CellNet classification scores for human ectoderm, endoderm, mesoderm, and germline cell types in control and *PRDM1*-KD cells at the DE stage. Bar plots show mean classification scores for the indicated cell types (*n* = 2 biological replicates). **(E)** Averaged plots of ChIP-seq signal centered on PRDM1 peaks for active (H3K27ac), active/primed enhancer-associated (H3K4me1), active/primed promoter-associated (H3K4me3), PRC2-mediated (H3K27me3), and PRC1-mediated (H2AK119ub1) marks in control and *PRDM1*-KD DE cells (*n* = 2 biological replicates). **(F)** Genome browser tracks showing ChIP-seq signal and RNA-seq coverage in control and *PRDM1*-KD DE cells at representative loci. **(G)** Heatmap showing the changes in ChIP-seq signal between *PRDM1*-KD and control cells at each individual PRDM1 peak at the DE stage, classified by ChromHMM categories. Gene ontology (GO) pathway enrichment of Biological Processes associated with each ChromHMM state was performed using GREAT (hg38) with default association rules. Statistical significance was determined using the binomial test, and multiple testing correction was applied using the Benjamini–Hochberg method (FDR ≤ 0.05). **(H)** Averaged plots of ChIP-seq signal for GATA6, SOX17, and EOMES TFs at PRDM1–pioneer TF co-bound sites, centered at PRDM1 peaks in control and *PRDM1*-KD cell lines at the DE stage. Bar plots showing log2 fold change in *PRDM1*, *GATA6*, *SOX17,* and *EOMES* mRNA expression between *PRDM1*-KD and control cells at the DE stage.

Next, to determine PRDM1’s function in epigenetic regulation of endoderm identity, we performed ChIP-seq for histone modifications enriched at PRDM1-bound sites (**Figure 1E**) – H3K27ac, H3K4me1, H3K4me3, H3K27me3, and H2AK119ub1 – in control and *PRDM1*-KD DE cells (**Figure 2E**). *PRDM1* knockdown resulted in a significant reduction in the Polycomb-associated modifications H2AK119ub1 and H3K27me3 at PRDM1-bound sites (both *P* ≤ 0.01), accompanied by an increase in the active histone mark H3K27ac (*P* ≤ 0.01) (**Figures 2E** and **S2E**). In contrast, no significant changes were observed in the active or primed marks H3K4me1 or H3K4me3 at PRDM1-bound sites (**Figures 2E** and **S2F**). Representative loci included the WNT signaling-associated gene *DACT1* and the liver-specific TF *PROX1*, both of which showed reduced Polycomb-associated modifications and de-repression upon *PRDM1* knockdown (**Figure 2F**). Although *PRDM1* knockdown consistently reduced H2AK119ub1 and H3K27me3 at PRDM1-bound sites across ChromHMM states, Polycomb-associated enhancers exhibited the most pronounced reductions (**Figure 2G**). Genes associated with Polycomb-associated enhancers were enriched for alternative-lineage-related biological processes such as “mesoderm formation”, “cardiac muscle differentiation”, and “neuron fate specification,” as well as for genes associated with tissues derived from the endoderm, including *HEY1, WNT5A, PAX6*, and *PROX1* (**Figure 2G**). These results suggest that in DE cells, PRDM1 contributes to the deposition and/or maintenance of Polycomb-associated modifications to repress alternative-lineage programs and prevent the precocious activation of genes associated with endoderm-derived tissues.

Lastly, to determine how PRDM1 binding influences pioneer TF occupancy, we performed ChIP-seq for EOMES, GATA6, and SOX17 in *PRDM1*-KD DE cells. *PRDM1* knockdown reduced, but did not abolish, the occupancy of all three pioneer TFs at PRDM1 co-bound sites (**Figure 2H** and **S2F**). Importantly, the expression of EOMES, GATA6, and SOX17 remained unchanged in *PRDM1-*KD cells (**Figure 2H**), indicating that the reduced occupancy was not a secondary effect of altered TF expression. While we previously showed that FOXA pioneer TFs are required for PRDM1 recruitment at the PFG stage,^35^ these findings unexpectedly indicate that PRDM1, in turn, contributes to stabilizing pioneer TF occupancy.

### The number of pioneer TF-binding sites determines the strength of PRDM1-mediated enhancer repression

We next aimed to define the direct functions and combinatorial regulatory logic of PRDM1 and the pioneer TFs (pTFs) EOMES, GATA6, and SOX17. We thus performed a lentivirus-based massively parallel reporter assay (lentiMPRA) on thousands of cis-regulatory sequences (CRSs) in DE cells, coupled with systematic TF-binding site (TFBS) mutations. To select TFBSs within CRSs, we developed a computational tool, MPRA-ChIP, which first identifies regions in which PRDM1 and pTF ChIP-seq peaks overlap within a 220-bp window, and then scans for their binding motifs using FIMO (Find Individual Motif Occurrences)^44^ (**Figure 3A**, steps 1-2). Among all PRDM1 ChIP-seq peaks in DE cells, 1,698 (48.7%) contained only a PRDM1 TFBS (PRDM1-only), whereas the remaining regions harbored a PRDM1 site together with EOMES, GATA6, and/or SOX17 TFBSs (PRDM1+pTF) (**Figure 3A**, step 2). For the lentiMPRA library, we first randomly downsampled the PRDM1-only CRS class to balance the number of sequences relative to PRDM1+pTF CRSs. We then generated a pool of CRSs consisting of wild-type (WT), TFBS mutation (MUT), and scrambled (SCRAM) negative-control sequences (**Figure 3A**, step 3). SCRAM sequences were generated by shuffling all nucleotides within each corresponding WT sequence. TFBS-mutated CRSs were generated in all possible combinations: with only the PRDM1 site mutated (PRDM1-only-MUT), with both the PRDM1 and pioneer TF sites mutated (PRDM1+pTF-MUT), or with only the pioneer TF sites mutated while keeping the PRDM1 site intact (pTF-MUT). All CRSs were cloned upstream of an SV40 promoter, followed by a unique 15-bp random barcode and an EGFP reporter in a lentiviral vector (**Figure 3A**, step 4). For robust quantification of transcriptional activity, we aimed to clone each CRS with 100 unique barcodes. We transduced the lentiMPRA library pool into hPSCs in four replicates using independent infections, performed differentiation into DE cells, and then sequenced genomic DNA and RNA (**Figure 3A**, step 4). To minimize bias from genomic integration sites, we targeted an average of 50 independent integration events per CRS–barcode pair. MPRAFlow^45^ was used for CRS– barcode association and barcode quantification, which showed high concordance across replicates for both RNA (average Pearson’s R = 0.96, *P* ≤ 0.01) and DNA (average Pearson’s R = 0.93, *P* ≤ 0.01) (**Figure S3A**). Each CRS was represented by a median of 62 DNA barcodes, and only 0.3% of CRSs lacked detectable barcodes (**Figure S3B**). To match the representation of WT and SCRAM sequences, we additionally removed three CRSs that were not detected in both groups. The final dataset therefore comprised 15,284 CRSs across WT, TFBS-MUT, and SCRAM groups (**Figure S3C**).

**Figure 3.**
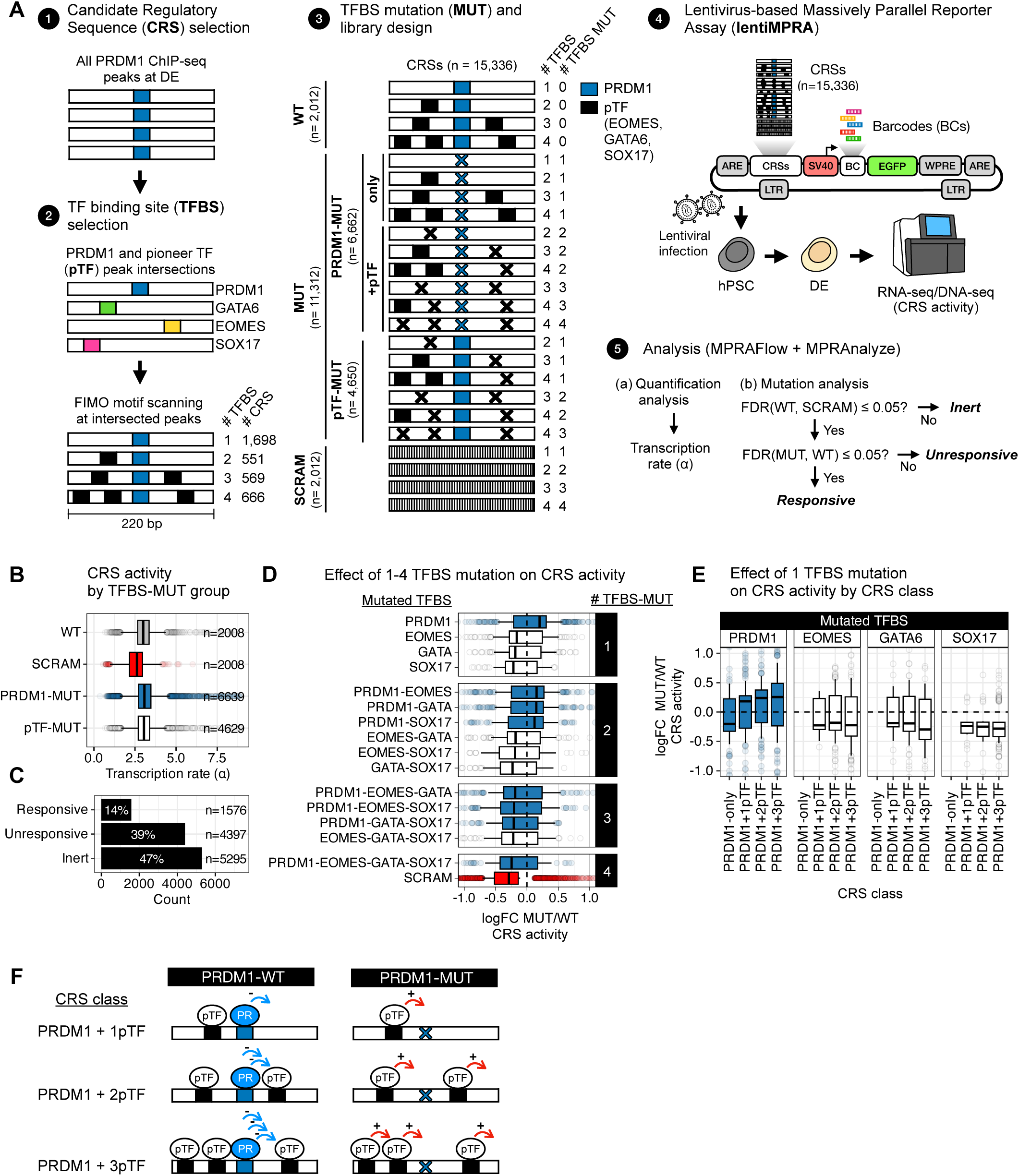
The number of pioneer TF-binding sites determines the strength of PRDM1-mediated enhancer repression. **(A)** Schematic representation of the lentiMPRA experimental design. Cis-regulatory sequence (CRS) candidates were derived from all PRDM1 ChIP-seq peaks identified at the DE stage. For TF-binding site (TFBS) selection, PRDM1 peaks were intersected with pioneer TFs (pTFs: GATA6, SOX17, and EOMES) peaks within a 220-bp window, followed by FIMO motif scanning. The library included wild-type (WT) and TFBS mutation (MUT) CRSs, as well as scrambled (SCRAM) sequences as negative controls. The selected CRSs and unique barcodes were cloned into a lentiviral vector containing the SV40 promoter and EGFP reporter. The infected hPSCs were differentiated to the DE stage, and DNA and RNA were extracted for sequencing (*n* = 4 independent infections). MPRAFlow was used for DNA and RNA barcode association and quantification. MPRAnalyze was used to estimate the transcriptional activity (α) driven by each CRS. **(B)** Box plots showing transcriptional activity (α) across WT, pTF-MUT, PRDM1-MUT (only and +pTF) CRSs, as well as SCRAM sequences. **(C)** Bar plots showing the proportion of CRSs in each activity category: *Inert, Unresponsive*, and *Responsive*. **(D)** Box plots showing fold changes in CRS activity between TFBS-MUT constructs and corresponding WT sequences across mutation groups. **(E)** Box plots showing fold changes in CRS activity between a single TFBS-MUT construct and corresponding WT sequences across CRS classes, stratified by the number of TFBSs. **(F)** Schematic model summarizing single TFBS-MUT results in lentiMPRAs.

To estimate CRS activity, we used MPRAnalyze^46^ to quantify the transcription rate (*a*) induced by each CRS (**Figure 3A**, step 5-a), with SCRAM sequences exhibiting the lowest *a* values overall (**Figure 3B**). To assess the regulatory function of TFBSs, we used the likelihood-ratio test implemented in MPRAnalyze (**Figure 3A**, step 5-b). We first classified WT CRS activity relative to SCRAM controls: CRSs whose activities were not significantly different from those of SCRAM sequences were classified as *Inert* (n = 5295, 47% of CRSs), whereas CRSs with significantly higher activities were classified as *Functional* (n = 5973, 53% of CRSs) (**Figure 3C**). *Functional* CRSs were then evaluated for sensitivity to TFBS mutation by comparing each mutant construct with its corresponding WT sequence. CRSs whose activities were not significantly altered by TFBS mutation were classified as *Unresponsive*, whereas those exhibiting significant changes were classified as *Responsive* (**Figure 3A**, step 5-b). Overall, 14% of all tested CRSs (n = 1576) were classified as *Responsive* to TFBS mutation and retained for downstream analyses (**Figure 3C**). The proportions of *Functional* CRSs and *Responsive* TFBSs were consistent with previous MPRA studies.^47–49^

Having identified *functional* CRSs whose activities in DE cells were affected by TFBS mutation, we evaluated the individual and combinatorial contributions of these TFBSs to CRS activity by comparing TFBS-MUT sequences with their corresponding WT sequences. Mutation of PRDM1 only (PRDM1-MUT) showed increased median CRS activity relative to WT, whereas mutation of a single pioneer TF site (EOMES, GATA6, or SOX17; 1 pTF) reduced median CRS activity (**Figure 3D** and **S3D**). These results indicate that PRDM1 represses CRS activity, whereas pioneer TFs promote CRS activity at these co-bound regulatory elements. Consistently, upon mutation of two TFBSs, PRDM1+pTF-MUT constructs showed increased median CRS activity, whereas mutations of two different pioneer TF sites (2pTF) showed reduced activity (**Figure 3D** and **S3D**). However, upon mutation of three or all four different TFBSs (PRDM1+2pTF, 3pTF, or PRDM1+3pTF), median CRS activity was generally reduced relative to WT, regardless of whether the PRDM1 motif was mutated or remained intact (**Figures 3D** and **S3D**). These results suggest that when only a single pioneer TF site remains intact, CRS activation is insufficient even in the absence of a functional PRDM1 site.

To quantify the effects of PRDM1 and pioneer TF site mutations, we modeled the log fold change (FC) as a function of the number of mutated TFBSs and mutation group (PRDM1-MUT or pTF-MUT) using a linear mixed model with CRS identity as a random effect. The model showed that CRS activity decreased significantly as the number of mutated TFBSs increased (*P* ≤ 0.001), while PRDM1 mutations resulted in higher CRS activity than pioneer TF mutations (*P* ≤ 0.001). To determine whether this overall group effect was maintained across CRS classes, we performed pairwise comparisons within each TFBS count class and observed higher activity for PRDM1-MUT than for pTF-MUT constructs at equivalent TFBS numbers (all *P* ≤ 0.01), indicating that CRS activity reflects a balance between PRDM1-mediated repression and pioneer TF-dependent activation (**Figure S3E**).

Finally, we assessed how a single TFBS mutation, targeting either PRDM1 or one pioneer TF, affected CRS activity across CRS classes stratified by TFBS number. Strikingly, the increase in CRS activity caused by PRDM1 TFBS mutation (PRDM1-MUT) became progressively greater as the number of different pioneer TF-binding sites increased (**Figure 3E** and **S3F**). In contrast, pioneer TFBS mutations (EOMES-MUT, GATA6-MUT, or SOX17-MUT) produced the opposite effect, leading to progressively larger reductions in CRS activity as the number of different pioneer TF-binding sites increased (**Figure 3E** and **S3F**). Altogether, these results suggest that the number of diverse pioneer TF-binding sites within a CRS determines the strength of its regulatory potential in both activating and repressive directions: pioneer TFs promote activation in the absence of PRDM1, whereas PRDM1 imposes increasingly strong repression at CRSs with greater pioneer TF co-occupancy (**Figure 3F**).

### Collaborative binding by diverse pioneer TFs and PRDM1 establishes focal chromatin accessibility and promotes hyper-bivalent enhancer formation

To define the mechanisms by which the number of diverse pioneer TF-binding sites determines the degree of PRDM1-mediated repression, we first assessed TF motifs and TFBS organization within CRSs, stratified by the number of co-bound pioneer TFs (EOMES, GATA6, and SOX17): PRDM1-only, PRDM1+1pTF, PRDM1+2pTF, and PRDM1+3pTF, including all possible combinations of different pioneer TFs (**Figure 3A**, step 2, and **Figure 4A**). Notably, the TF motif stringency score (reflecting similarity to the canonical TF motif) for PRDM1, but not for the pioneer TFs, decreased as the number of pioneer TF-binding sites increased (**Figure 4B**), suggesting that PRDM1 occupies increasingly suboptimal binding sites in the context of greater pioneer TF co-binding. Previous studies demonstrate that suboptimal, low-affinity motifs can confer specificity on enhancer activity and facilitate cooperative and collaborative TF binding.^50–52^ The total span of the motif module increased with TFBS content within a CRS, from ∼50 bp in PRDM1+1pTF CRSs to ∼100 bp in PRDM1+3pTF CRSs, while remaining within the 147-bp DNA footprint of a single nucleosome (**Figure 4C**, left). Within CRSs, the average pairwise distance between TF motifs ranged from approximately 48 to 58 bp (**Figure 4C**, right). These results align with previous studies on gene activation, showing that TF cooperation is enhanced between TFBSs spaced approximately 50 bp apart, which can promote nucleosome-mediated TF cooperativity.^2,3^

**Figure 4.**
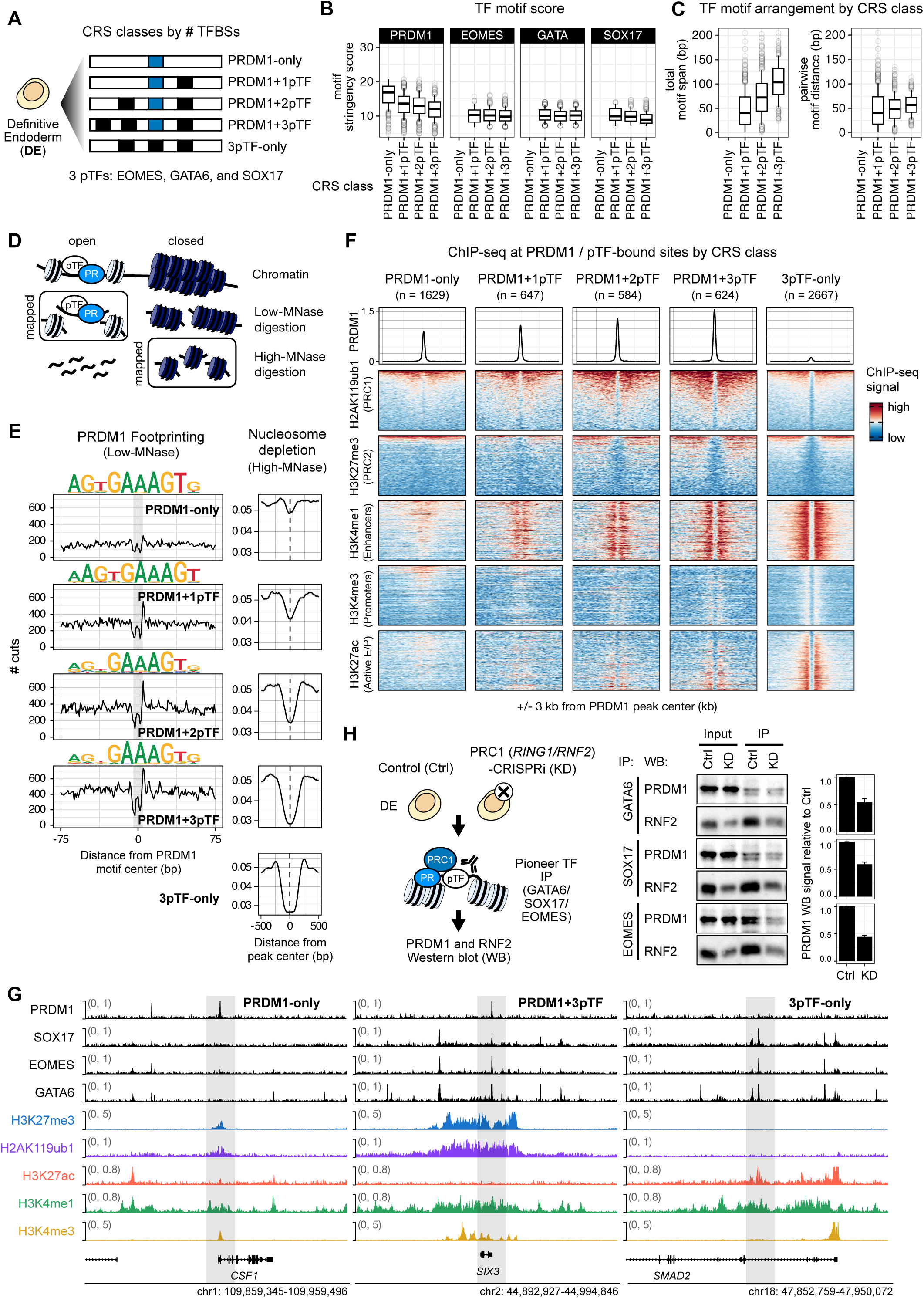
Collaborative binding by diverse pioneer TFs and PRDM1 establishes focal chromatin accessibility and promotes hyper-bivalent enhancer formation. **(A)** Schematic representation of cis-regulatory sequence (CRS) classification based on TF-binding site (TFBS) content at the DE stage. **(B)** Box plots of motif stringency scores for PRDM1, EOMES, GATA6, and SOX17 TFBSs across CRS classes. **(C)** Box plots of the mean pairwise motif distance and total motif span across CRS classes. **(D)** Schematic representation of Low- and High-MNase-seq experiments. **(E)** Aggregated PRDM1 footprinting (Low-MNase-seq) and nucleosome occupancy profiles (High-MNase-seq) at PRDM1 and/or pioneer TF-bound sites in DE cells. Above the Low-MNase profiles, the top PRDM1 HOMER motifs identified within each CRS class are shown. **(F)** Averaged PRDM1 ChIP-seq profiles and heatmaps of ChIP-seq signal for histone modifications (H2AK119ub1, H3K27me3, H3K4me1, H3K4me3, and H3K27ac) centered at PRDM1 peaks or 3pTF-only peaks across CRS classes. **(G)** Genome browser tracks showing TF and histone modification ChIP-seq signal across CRS classes. **(H)** Schematic representation of PRDM1 and pTFs co-immunoprecipitation (co-IP) experiments in control and PRC1 (*RING1/RNF2*)-CRISPRi (KD) cell lines in DE cells. IP of pioneer TFs followed by Western blot (WB) for PRDM1 and RNF2 in control and *PRC1*-KD cells at the DE stage (middle). Quantification of Western blot signal for PRDM1 upon IP of GATA6, SOX17, and EOMES in control and *PRC1*-KD cells (right).

Next, to investigate nucleosome organization at PRDM1-bound sites with varying numbers of pioneer TF-binding sites, we performed Low- and High-MNase-seq in DE cells to profile subnucleosomal TF-protected footprints under Low-MNase conditions, as well as stable nucleosomes under High-MNase conditions (**Figures 4D** and **S4A**).^53,54^ Footprinting analysis revealed progressively stronger protection centered on PRDM1 motifs as the number of co-bound pioneer TFs increased, suggesting increased stability of PRDM1 occupancy (**Figure 4E**). *De novo* motif analysis using HOMER^55^ further showed that while the common GAAGT core of the PRDM1 motif was maintained across all classes, the 5’ and 3’ flanking positions became increasingly degenerate as the number of co-bound pioneer TF increased (**Figure 4E**), consistent with the TF motif stringency analysis (**Figure 4B**). Furthermore, High-MNase-seq analysis showed progressively greater nucleosome depletion around PRDM1-bound sites with increasing pioneer TF co-binding (**Figure 4E**). Notably, the extent of nucleosome depletion at PRDM1+3pTF sites was comparable to that at 3pTF-only sites, while exhibiting slightly narrower depletion (**Figure 4E**). PRDM1 ChIP-seq signal also exhibited progressively stronger peaks as the number of co-bound pioneer TFs increased (**Figure 4F**, top). Together, these results suggest that collaborative binding by diverse pioneer TFs promotes focal nucleosome remodeling and stabilizes PRDM1 occupancy in a pioneer TF dose-dependent manner.

We next examined how focal chromatin accessibility and increased PRDM1 occupancy were translated into local epigenetic states. Strikingly, enrichment of the bivalent enhancer-associated modifications H2AK119ub1, H3K27me3, and H3K4me1 progressively increased with the number of pioneer TFs co-bound with PRDM1, culminating in the formation of “hyper-bivalent” enhancers at highly occupied PRDM1+3pTF sites (**Figure 4F**). Consistent with this, occupancy of the PRC1 subunits RNF2 and RYBP and the PRC2 subunit SUZ12 also increased with the number of pioneer TFs co-bound with PRDM1 (**Figure S4B**). These enhancers showed approximately 4- to 5-fold higher H2AK119ub1 and H3K27me3 signals than PRDM1-only CRSs, with median peak lengths approximately 2-fold longer (**Figure S4C**). Importantly, Polycomb-associated modifications were not enriched at 3pTF-only sites. Instead, these regions were strongly enriched for the active enhancer-associated histone modifications H3K27ac and H3K4me1 (**Figure 4F**), indicating that PRDM1 co-occupancy distinguishes hyper-bivalent enhancers from active enhancers bound by the same three pioneer TFs. Representative loci include *CSF1*, a regulator of macrophage development and survival, at a PRDM1-only locus; *SIX3*, a TF involved in forebrain development, at a PRDM1+3pTF locus; and *SMAD2*, a mediator of TGFβ/NODAL signaling, at a 3pTF-only locus (**Figure 4G**). Together, these results show that collaborative binding by diverse pioneer TFs amplifies PRDM1-mediated Polycomb epigenetic states, culminating in the establishment of hyper-bivalent enhancers at highly occupied loci.

To further investigate the molecular interactions underlying collaborative binding by pioneer TFs and PRDM1, we used a Dox-inducible CRISPRi cell line targeting the PRC1 core subunits *RING1* and *RNF2* (*PRC1-*KD),^35^ and assessed the impact of *PRC1* knockdown on interactions between PRDM1 and pioneer TFs in DE cells (**Figure 4H**). We first confirmed that *PRC1* knockdown did not alter PRDM1 or pioneer TF expression levels (**Figure S4D**). Co-IP experiments revealed that PRDM1 physically interacted with EOMES, GATA6, and SOX17 in control DE cells and that these interactions were reduced by 40–60% upon *PRC1* knockdown (**Figure 4H**). Together with PRDM1–pTF-associated PRC enrichment (**Figure 1H**, **1I**, **2E**) and PRDM1’s contribution to pTF occupancy (**Figure 2H**), these findings suggest that PRDM1, pioneer TFs, and PRC1 mutually reinforce their interactions, contributing to stable and collaborative occupancy.

### Hyper-bivalent enhancers reinforce the repression of alternative-lineage, past and future developmental programs

To define the functional properties of hyper-bivalent enhancers established by diverse pioneer TFs (EOMES, GATA6, and SOX17) and PRDM1, we first annotated each CRS to its nearest gene using the Genomic Regions Enrichment of Annotations Tool (GREAT).^56^ We found that CRSs containing a greater number of pioneer TF-binding sites were located progressively farther from transcription start sites (TSSs), suggesting that they regulate genes across greater genomic distances (**Figure 5A**).

**Figure 5.**
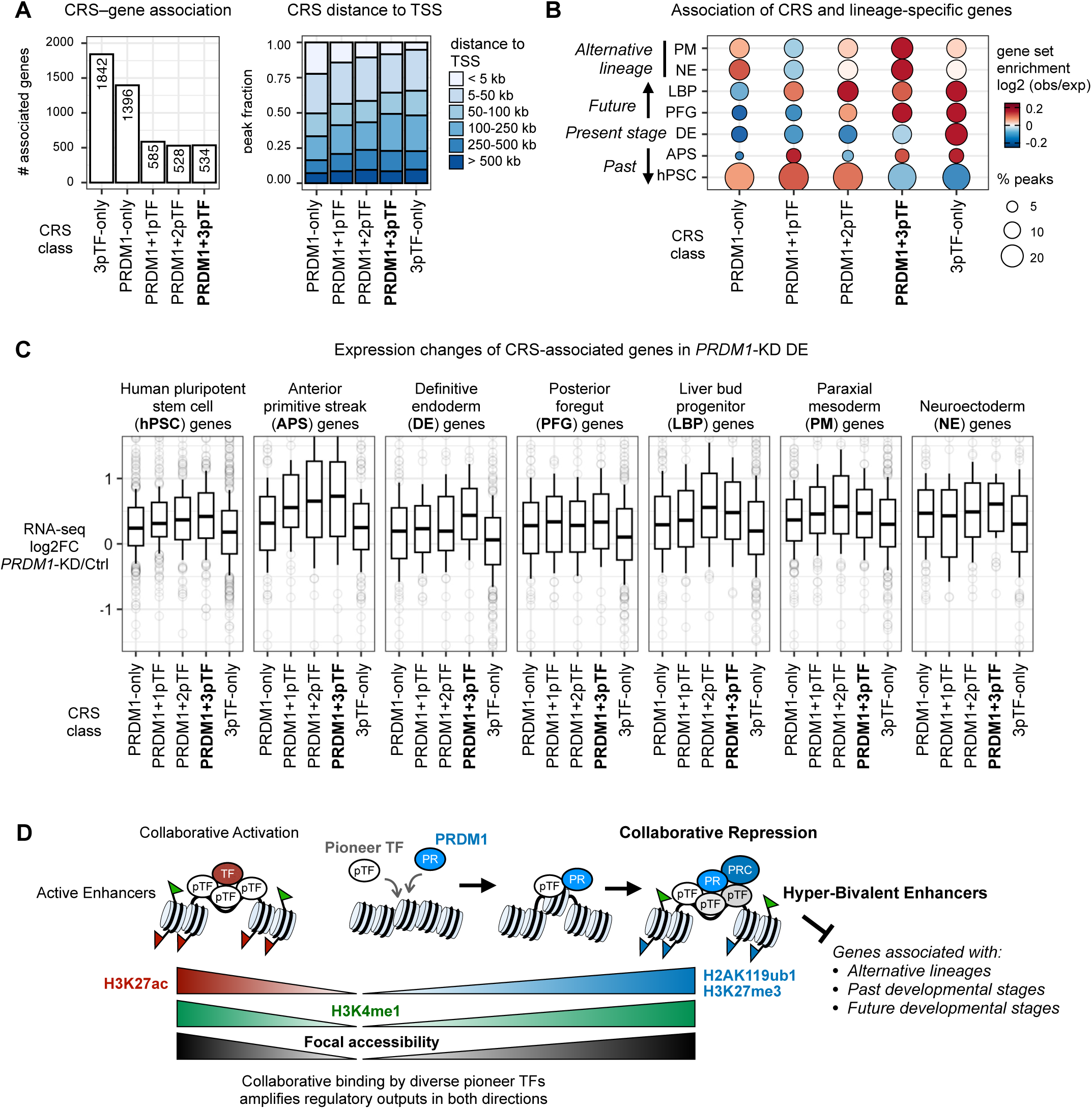
Hyper-bivalent enhancers reinforce the repression of alternative-lineage, past and future developmental programs. **(A)** Bar plots showing the number of genes associated with each cis-regulatory sequence (CRS) class using the nearest gene association rule of the GREAT annotation software (left panel). Stacked bar plots showing the distribution of distances between peaks and transcription start sites (TSSs) across CRS classes (right panel). **(B)** Dot plot showing enrichment of lineage-specific gene programs across CRS classes. The size of the dots represents the proportion of lineage-specific genes enriched in each CRS class. Log2 (obs/exp) stands for the base-2 logarithm of the observed-to-expected ratio. **(C)** Box plots of expression changes of lineage-specific genes upon *PRDM1*-CRISPRi knockdown at the DE stage across CRS classes. **(D)** Schematic model summarizing a “collaborative repression” model in which collaborative binding by diverse pioneer TFs reinforces the context-specific assembly of repressive regulators to safeguard lineage and developmental fidelity. This model redefines the conventional view that pioneer TF-mediated accessible chromatin primarily serves as a platform for enhancer activation and instead supports that focal accessibility can amplify regulatory outputs in both directions – activation and repression.

We then tested whether PRDM1-only, PRDM1+pTF, and 3pTF-only CRSs were associated with distinct lineage- and stage-specific developmental gene programs. Using RNA-seq data,^35^ we defined gene sets specific to alternative lineages (PM and NE), as well as past (hPSC and APS), present (DE), and future (PFG and LBP) endodermal stages. Notably, PRDM1+3pTF CRSs were strongly enriched near genes specific to alternative lineages (PM and NE), as well as past (APS) and future (PFG and LBP) endoderm developmental stages (**Figure 5B**). Importantly, PRDM1+3pTF CRSs were not enriched near the present-stage (DE) genes. In contrast, the 3pTF-only CRSs were strongly enriched near endoderm-lineage genes, including DE-specific genes, but not near alternative-lineage gene sets (**Figure 5B**). These results suggest that hyper-bivalent enhancers regulate alternative-lineage, and past and future developmental programs. We next assessed the effects of *PRDM1*-CRISPRi knockdown on the expression of genes associated with these CRS classes. Notably, *PRDM1* knockdown led to progressively greater de-repression of genes associated with PRDM1+pTF CRSs as the number of co-bound pioneer TFs increased (**Figure 5C**). These results are consistent with the effects of PRDM1 TFBS mutation on CRS activity observed in the lentiMPRA (**Figure 3E**). As expected, *PRDM1* knockdown did not markedly alter the expression of genes associated with 3pTF-only CRSs (**Figure 5C**). Together, these results suggest that hyper-bivalent enhancers reinforce developmental fidelity and robustness during endoderm differentiation by preventing alternative-lineage differentiation, restraining reactivation of earlier developmental programs, and preventing precocious activation of later programs.

## DISCUSSION

In this study, we address the fundamental gap of how broadly expressed repressive TFs and epigenetic regulators achieve context-specific repression. Our results demonstrate that the repressive TF PRDM1 cooperates with lineage-and stage-specific combinations of pioneer TFs during endoderm differentiation. Furthermore, the diverse pioneer TFs EOMES, GATA6, and SOX17 synergistically promote local nucleosome remodeling and stabilize PRDM1 occupancy, thereby enhancing PRC recruitment and establishing hyper-bivalent enhancers that repress alternative lineage, past and future developmental programs. These findings challenge the activation-centric view of TF cooperation and support a “collaborative repression” model in which collaborative binding by diverse pioneer TFs reinforces the context-specific assembly of repressive regulators to safeguard lineage fidelity (**Figure 5D**). This model is complementary to, yet distinct from, previously proposed mechanisms of pioneer-TF-mediated repression^57^, which include enhancer reorganization and decommissioning,^58–60^ TFBS competition,^61^ and “pioneer repression” in which BACH1 regulates chromatin opening and repression.^62^ Our model challenges the prevailing view that pioneer TF-mediated accessible chromatin primarily serves as a platform for enhancer activation and instead supports that focal accessibility can amplify regulatory outputs in both directions – activation and repression (**Figure 5D**).

Hyper-bivalent enhancers share conceptual features with super-enhancers,^5,6^ including collaborative TF binding and amplified regulatory output in directing cell identity, but they instead function as repressive regulatory elements. Clusters of suboptimal, low-affinity TF motifs are also increasingly recognized as a mechanism that promotes cell-type specificity and regulatory robustness at developmental enhancers.^50–52,63^ Our data show that PRDM1 motifs become progressively more degenerate at sites with greater pioneer TF co-occupancy, suggesting that collaborative binding at suboptimal motifs can provide regulatory specificity not only for transcriptional activation but also for context-specific repression.

We demonstrate that PRDM1 dynamically shifts its genomic occupancy from definitive endoderm to foregut endoderm by cooperating with distinct lineage- and stage-specific combinations of pioneer TFs: EOMES, GATA6, and SOX17 in DE; FOXA TFs,^35^ likely with other pioneer TFs (e.g., GATA and SOX4) in PFG. Intriguingly, human primordial germ cell (PGC) specification also depends on SOX17 and PRDM1, which contribute to repression of endodermal and mesodermal programs and activation of the germ cell program.^34,64–66^ In this context, however, SOX17 and PRDM1 ChIP-seq peaks show minimal overlap: SOX17 predominantly occupies distal regulatory regions and primarily functions as an activator, whereas PRDM1 exhibits pronounced promoter binding and primarily functions as a repressor.^67^ Primordial germ cells do not express EOMES and GATA6; instead, they express other pioneer TFs, such as OCT4 and TFAP2C, which co-localize with PRDM1.^66–68^ These observations suggest that PRDM1 may cooperate with a distinct, PGC-specific combination of pioneer TFs to repress inappropriate somatic programs. We hypothesize that context-specific collaborative TF binding is determined by the availability of TFs in a given cellular state together with the genomic distribution of their cognate TFBSs.

In *Drosophila*, PRCs are recruited to specific regulatory elements known as Polycomb response elements, which are frequently co-occupied by Trithorax group proteins to form bivalent epigenetic states. Because Polycomb regulates a similar set of developmental genes in *Drosophila* and vertebrates,^69^ the field has sought to identify vertebrate counterparts based on sequence conservation, but with limited success.^30,31,69,70^ Intriguingly, human hyper-bivalent enhancers share several conceptually analogous features with *Drosophila* Polycomb response elements: (1) flexible clusters of multiple TF motifs,^71^ (2) nuclease-hypersensitive (accessible) chromatin occupied by the pioneer TF GAF,^72–74^ and (3) PRC1-dependent stabilization of TF binding at suboptimal motifs.^75^ These similarities suggest that, even though they employ distinct TFs, *Drosophila* Polycomb response elements and human hyper-bivalent enhancers share a common regulatory logic.

The concept of pioneer TFs emerged from efforts to understand how TFs initiate chromatin opening and potentiate transcriptional activation during development within naïve or silent chromatin.^15,76^ Our study expands this paradigm by demonstrating that pioneer TF-mediated chromatin accessibility can also facilitate the establishment of repressive states. This principle may also have implications for pioneer TF-induced direct cellular reprogramming, where achieving high-fidelity cell identities requires not only target gene activation but also effective repression of inappropriate programs. More broadly, our findings support a principle in which collaborative binding by diverse pioneer TFs confers context specificity on broadly expressed repressive regulators, thereby safeguarding lineage and developmental fidelity during development.

## ACKNOWLEDGMENTS

We thank Ken Zaret, Aaron Zorn, Kohta Ikegami, and Brian Gebelein for helpful comments; Natascha Bushati for editing the paper; and the Digestive Disease Research Core Center at CCHMC (Pluripotent Stem Cell Facility, DNA sequencing Core, Bio-Imaging and Analysis Facility). This work was supported by the National Institutes of Health [P30 DK078392 and R01GM143161 to M.I.]; Cincinnati Children’s Research Foundation [Trustee Award to M.I. and H-W.L., Center for Pediatric Genomics Pilot Award to M.I.]

## AUTHOR CONTRIBUTIONS

Conceptualization: M.I. and G.M.; investigation: G.M., M.B., K.L., S.M., S.S.; formal analysis: G.M., H-W.L.; software: H-W.L., G.M.; visualization: G.M., M.I.; writing – original draft: M.I., G.M.; writing – review & editing: all authors; supervision and funding acquisition: M.I. and H-W.L.

## KEY RESOURCES TABLE

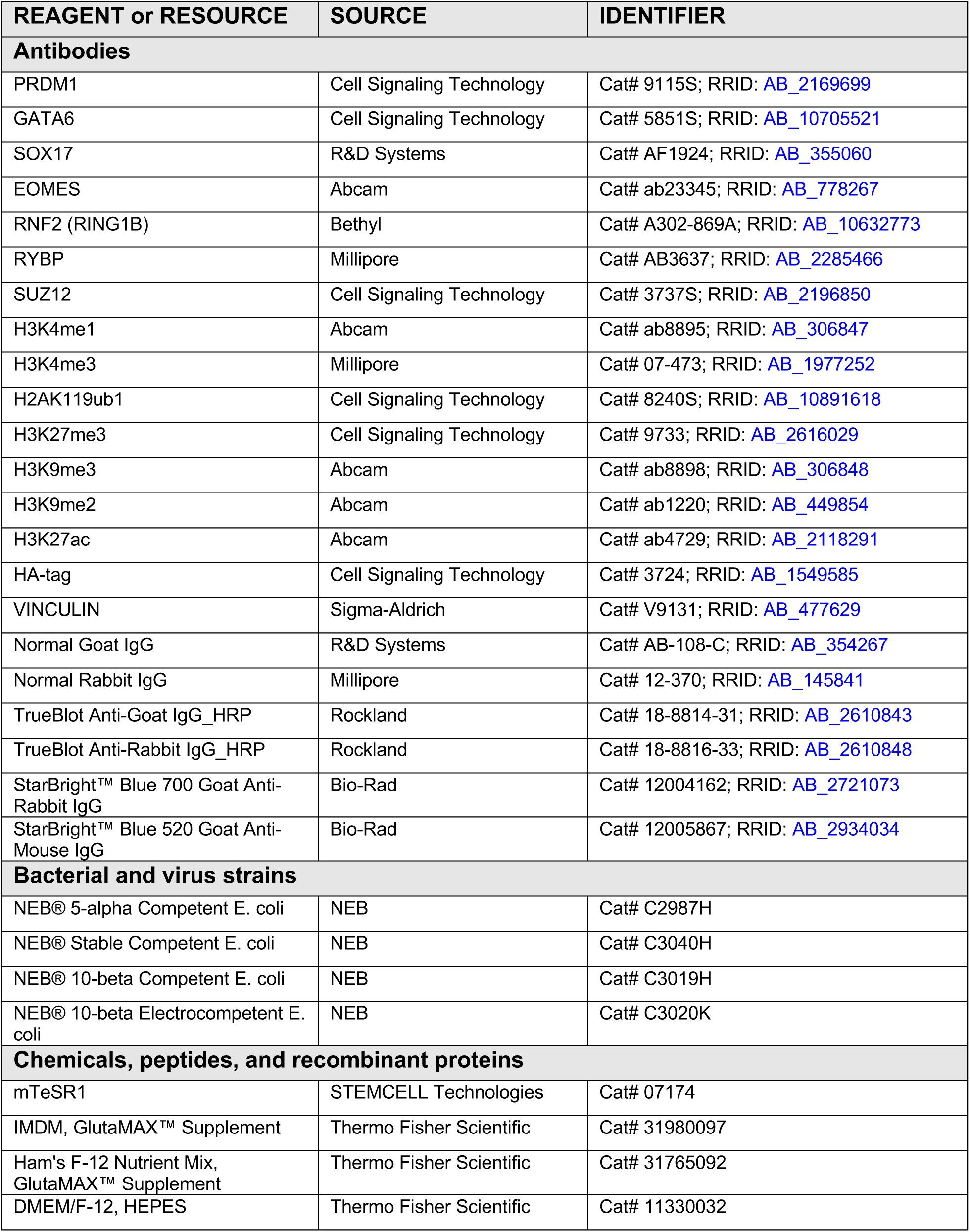
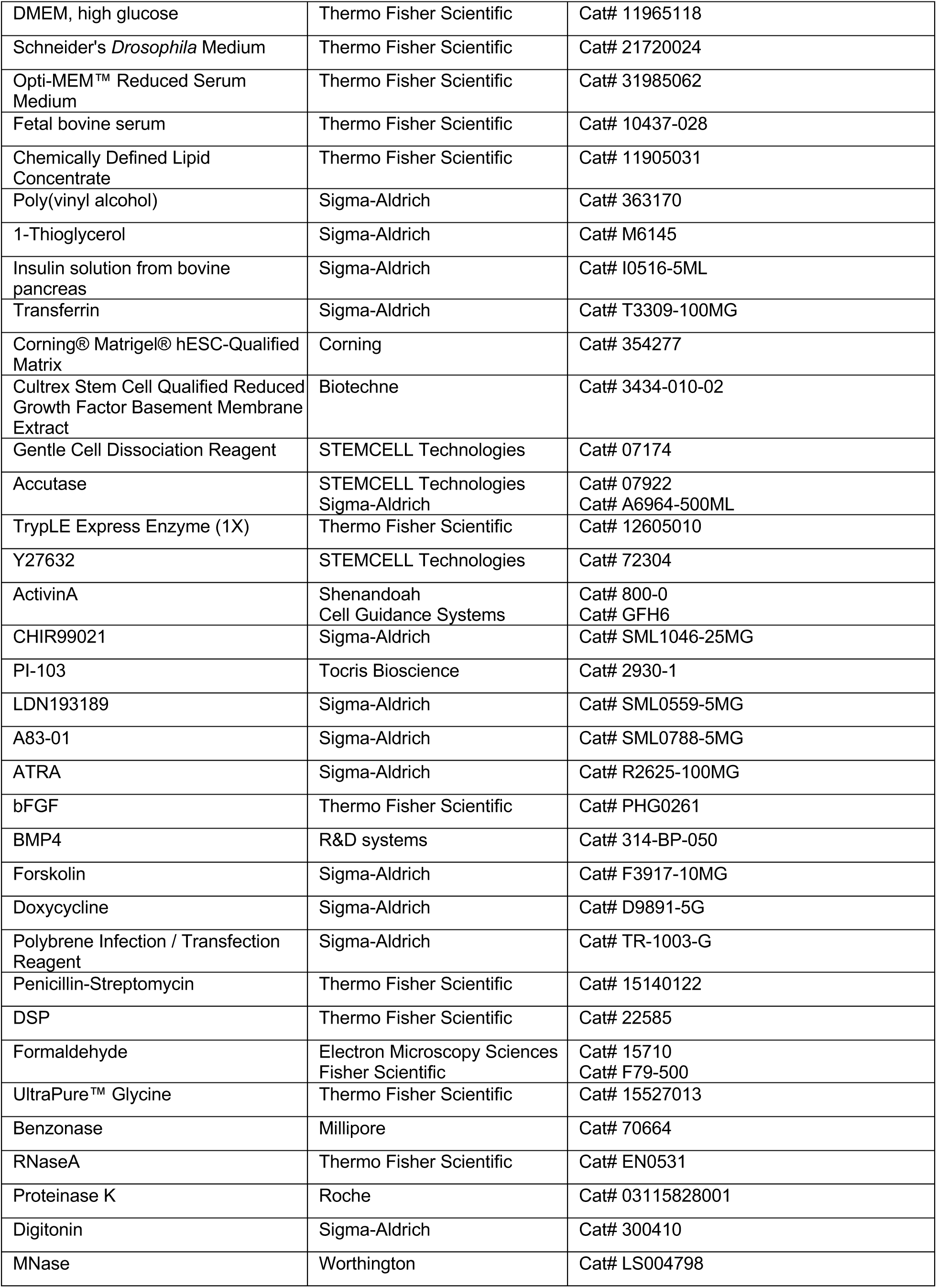
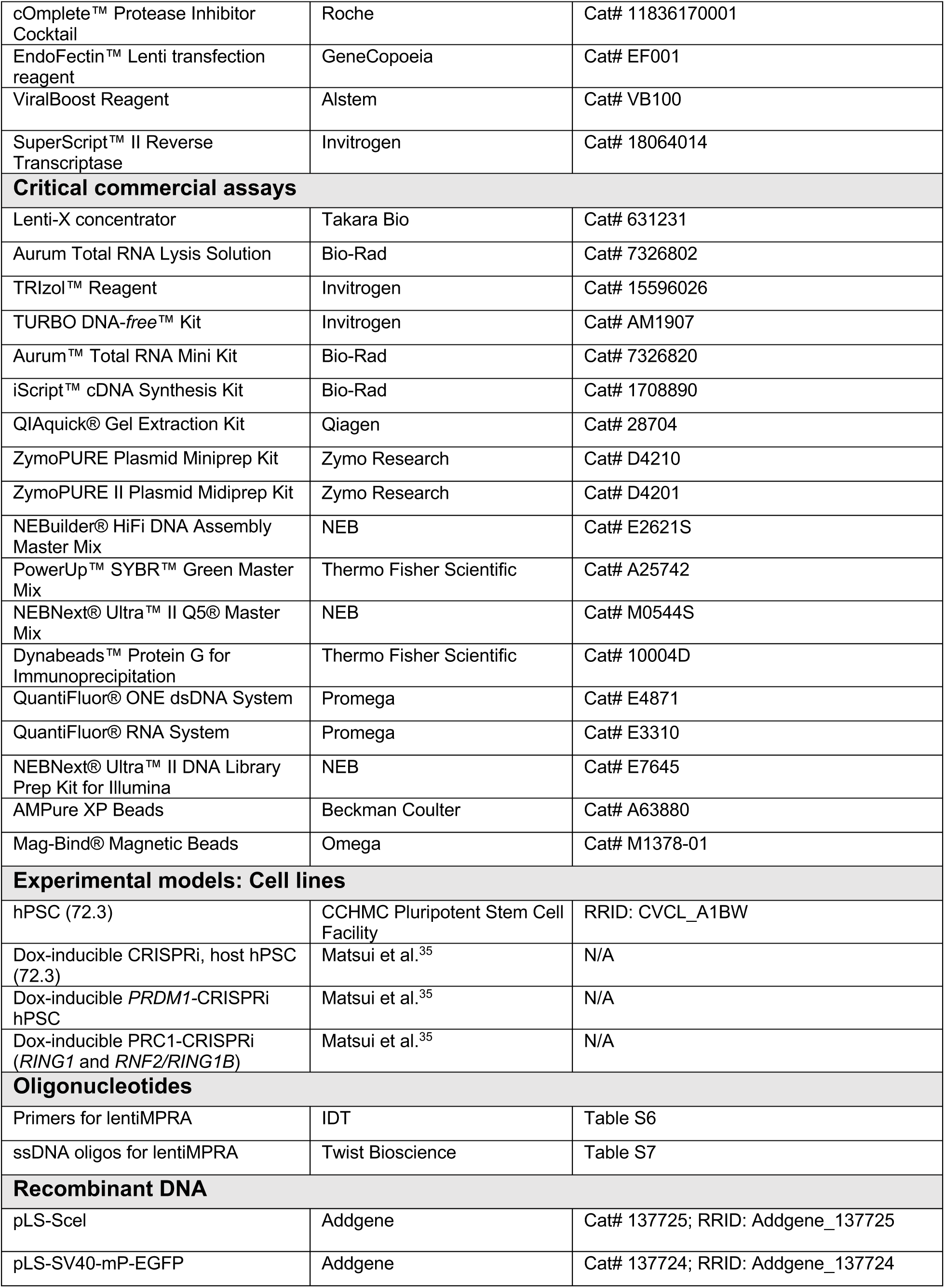
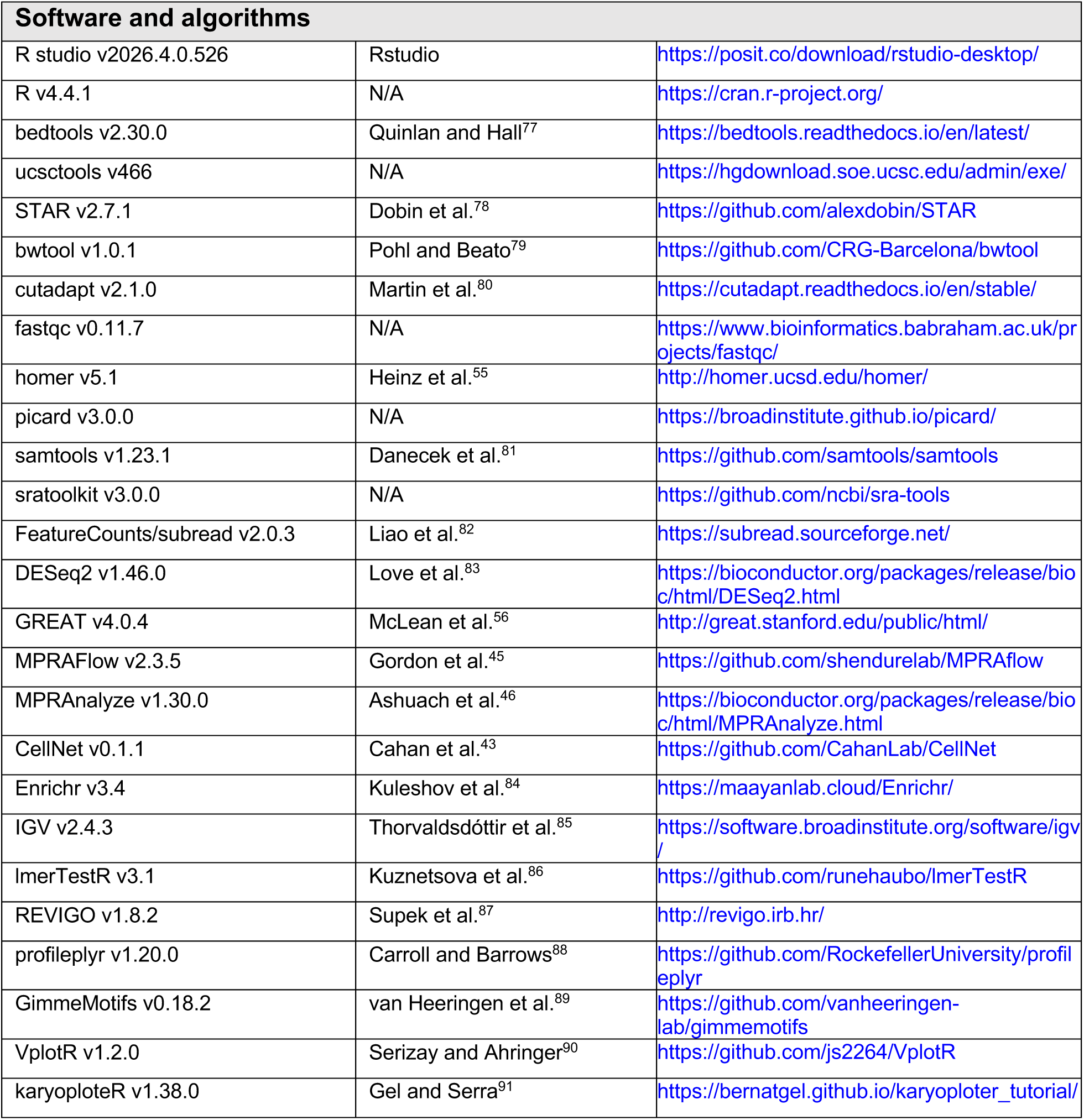

## EXPERIMENTAL MODEL AND STUDY PARTICIPANT DETAILS

### Culture of human induced pluripotent stem cells

Human induced pluripotent stem cells (hPSCs; line 72.3, male, RRID: CVCL_A1BW) were obtained from the CCHMC Pluripotent Stem Cell Facility (PSCF)^92^ and cultured under feeder-free conditions in mTeSR1 (85850, STEMCELL Technologies) on plates coated with Matrigel (354277, Corning) or Cultrex Stem Cell Qualified Reduced Growth Factor Basement Membrane Extract (3434-010-02, Bio-Techne). All hPSCs were maintained in a humidified incubator at 37 °C with 5% CO₂. Spontaneously differentiated cells were manually removed prior to passaging and differentiation. For passaging, hPSC colonies were dissociated into small cell clusters using Gentle Cell Dissociation Reagent (07174, STEMCELL Technologies) and seeded onto Matrigel- or Cultrex-coated plates in mTeSR1.^93^

Dox-inducible CRISPRi host hPSCs were distributed by the PSCF from a quality-controlled, cryopreserved bank prepared at passage 50, transduced with gRNAs and used in experiments up to passage 80 (*PRDM1-*CRISPRi)^35^ and passage 73 (*PRC1-*CRISPRi)^35^. Cells were confirmed to be mycoplasma-free, to harbor a normal G-banded karyotype, and to have verified identity by STR profiling. To induce CRISPRi-mediated knockdown of target genes, cells were cultured in the presence of 2 µM doxycycline (D9891-5G, Sigma-Aldrich). CRISPRi hPSC lines are listed in the **Key Resources Table**.

## METHOD DETAILS

### hPSC to endoderm differentiation

For cell differentiation, hPSCs were dissociated into single cells using Accutase (07922, STEMCELL Technologies; A6964-500ML, Sigma-Aldrich). Dissociated cells were plated onto Matrigel- or Cultrex-coated culture plates and cultured in mTeSR1 supplemented with 10 µM Y-27632 (72304, STEMCELL Technologies) one day prior to differentiation. Chemically defined basal media 2 (CDM2) and 3 (CDM3) were prepared as previously reported,^94^ filtered through a 0.22-µm filter unit, and stored at -80 °C until use. Once thawed, CDM2 and CDM3 were used within 2 – 3 weeks.

Endodermal induction was performed as previously described.^35,37,94^ For anterior primitive streak (APS) induction, hPSCs were cultured in CDM2 supplemented with 100 ng/mL Activin A (800-0, Shenandoah; or GFH6, Cell Guidance Systems), 2 µM CHIR99021 (SML1046-25MG, Sigma-Aldrich), and 50 nM PI103 (2930-1, Tocris Bioscience) for 24 h. For definitive endoderm (DE) induction, APS cells were briefly washed with DMEM/F12 and cultured in CDM2 supplemented with 100 ng/mL Activin A and 250 nM LDN193189 (SML0559-5MG, Sigma-Aldrich) for 24 h. For posterior foregut (PFG) induction, DE cells were briefly washed with DMEM/F12 and cultured in CDM3 supplemented with 1 µM A83-01 (SML0788-5MG, Sigma-Aldrich), 2 µM all-trans retinoic acid (ATRA; R2625-100MG, Sigma-Aldrich), 10 ng/mL bFGF (PHG0261, Thermo Fisher Scientific), and 30 ng/mL BMP4 (314-BP-050, R&D Systems) for 24 h. For liver bud progenitor (LBP) induction, PFG cells were cultured in CDM3 supplemented with 10 ng/mL Activin A, 30 ng/mL BMP4, and 1 mM forskolin (F3917-10MG, Sigma-Aldrich) for 72 h. Differentiation media was changed every 24 h.

### Western blotting

Between 1 and 2.5 µg of nuclear protein in Laemmli Sample Buffer (J60015, Alfa Aesar) was denatured at 98 °C for 5 min, resolved on 10%, 12.5%, or 15% SDS-PAGE gels at 60 V for 30 min followed by 100 V for 2 h, and transferred onto either 0.2- or 0.45-µm pore-size PVDF membranes (for chemiluminescence; LC2002, Thermo Fisher Scientific; IPSN07852, Millipore) or 0.2-µm nitrocellulose membranes (for fluorescence; 1620112, Bio-Rad) using a wet tank transfer system with a Mini Blot Module (NW2000, Thermo Fisher Scientific) at 20 V (chemiluminescence) or 10 V (fluorescence) for 1 h.

Transferred membranes were briefly washed with TBST (0.1% Tween-20 in TBS, for chemiluminescence) or TTBS (0.05% Tween-20 in TBS, for fluorescence) and blocked for 1 h at room temperature in blocking buffer consisting of either 5% Blotting-Grade Blocker (1706404, Bio-Rad) or 5% BSA (A9647-50G, Sigma-Aldrich) prepared in TBST (chemiluminescence) or TBS (fluorescence). Membranes were then washed with TBST or TTBS and incubated with primary antibodies diluted in blocking buffer at 4 °C overnight. The following day, membranes were washed three to five times with TBST or TTBS for 5 min at room temperature and incubated with secondary antibodies diluted in 1% BSA in TBST (chemiluminescence) or 5% Blotting-Grade Blocker in TBS (fluorescence) for 1 h at room temperature. Membranes were washed three to five additional times with TBST or TTBS for 5 min at room temperature. Chemiluminescent signals were developed using ECL Select Western Blotting Detection Reagent (RPN2235, GE Healthcare) or Clarity™ Western ECL Substrate (170-5061, Bio-Rad) for 5 min. Chemiluminescence and fluorescence images were acquired using a ChemiDoc™ MP Imaging System (12003153, Bio-Rad). Antibodies used in the Western blot assays are listed in **Table S3**.

### Co-immunoprecipitation

Co-immunoprecipitation (co-IP) was performed with modifications to previously published protocols.^20,95^ Cultured cells were briefly washed with DMEM/F12 and crosslinked with 1.5 mM DSP (22585, Thermo Fisher Scientific) in 1x PBS at room temperature for 30 min with gentle shaking. The crosslinking solution was aspirated, and the reaction was quenched with 30 mM Tris-HCl (pH 7.4) in 1x PBS for 20 min with gentle shaking. Crosslinked cells were washed twice with ice-cold 1x PBS and scraped from the plate in ice-cold 0.01% PVA-supplemented 1x PBS. The cell suspension was transferred to conical tubes and centrifuged at 350 xg for 3-5 min at 4°C. The supernatant was discarded, and the cell pellet was resuspended in ice-cold 0.01% PVA-supplemented 1x PBS and centrifuged at 800 xg for 3-5 min at 4 °C. The supernatant was discarded, and the cell pellet was snap-frozen on dry ice and stored at -80 °C until further processing.

For nuclear protein extraction, frozen cell pellets were resuspended in Cell Lysis Buffer (5 mM HEPES-NaOH [pH 7.9], 10 mM KCl, 1 mM DTT, 0.5% NP-40, 1x protease inhibitor) and incubated on ice for 10 min. The lysate was centrifuged at 1700 xg for 10 min at 4 °C, and the supernatant was discarded. The nuclear pellet was resuspended in Nuclear Lysis Buffer (100 mM NaCl, 25 mM HEPES-NaOH [pH 7.9], 1 mM MgCl₂, 0.2 mM EDTA, 0.5% NP-40, 600 U/mL Benzonase [70664, Millipore], 1x protease inhibitor) and incubated for 4 h at 4 °C with rotation. The NaCl concentration of the nuclear lysate was adjusted to 200 mM and incubated for an additional 30 min at 4 °C with rotation. The nuclear lysate was centrifuged at maximum speed (16000 xg) for 30 min at 4 °C. A small volume of the precleared lysate was reserved as Input, and the remaining lysate was incubated with the appropriate amount of primary antibody (**Table S4**) overnight at 4 °C with rotation.

For each immunoprecipitation reaction, 50 µL Dynabeads Protein G (10004D, Thermo Fisher Scientific) were washed twice with PBST (0.1% Tween-20 in PBS) and incubated with the antibody-lysate mixture for 2 h at 4 °C with rotation. Dynabeads were washed five times with IP Wash Buffer (100 mM NaCl, 25 mM HEPES-NaOH [pH 7.9], 1 mM MgCl₂, 0.2 mM EDTA, 0.5% NP-40, 1x protease inhibitor), with each wash performed for 2 min at room temperature with rotation. Dynabead-bound protein complexes (IP samples) were eluted in 1x Laemmli Sample Buffer (1610737EDU, Bio-Rad) for 20 min at 65 °C with shaking at 1000 rpm in a ThermoMixer F1.5 (EP5384000020, Eppendorf). For denaturing IP and Input samples, 1/20 volume of 2-mercaptoethanol was added, followed by boiling for 5 min at 98 °C. Denatured samples were stored at -20 °C until analysis by Western blotting. Antibody information is listed in **Table S4**.

### Chromatin immunoprecipitation

Cells were crosslinked with 1% formaldehyde (F79-500, Fisher Scientific) in 1x PBS for 10 min at room temperature. The crosslinking reaction was then quenched by adding 0.125 M glycine (15527013, Thermo Fisher Scientific) for 5 min at room temperature. Crosslinked cells were centrifuged at 600 xg for 5 min at 4 °C, and the supernatant was discarded. The cell pellet was washed twice with ice-cold 1x PBS, centrifuged at 600 xg for 5 min at 4 °C, snap-frozen on dry ice, and stored at -80 °C.

Crosslinked cells were resuspended in Lysis Buffer 1 (10 mM Tris-HCl [pH 8.0], 10 mM NaCl, 0.5% NP-40, and 1x Complete Protease Inhibitor, EDTA-free) and incubated on ice for 10 min. The cells were then centrifuged at 665 xg for 5 min at 4 °C, and the supernatant was discarded. The cell pellet was resuspended in Lysis Buffer 2 (50 mM Tris-HCl [pH 8.0], 10 mM EDTA, 0.32% SDS, and 1x Complete Protease Inhibitor, EDTA-free) and incubated on ice for 10 min. The lysate was diluted with IP Dilution Buffer (20 mM Tris-HCl [pH 8.0], 150 mM NaCl, 2 mM EDTA, 1% Triton X-100, and 1x Complete Protease Inhibitor, EDTA-free) and transferred to a milliTUBE ATA Fiber (520135, Covaris). Chromatin was sonicated using a Covaris S220 (500217, Covaris) for 3.5 min (for TF ChIP) or 5.5 min (for histone ChIP). Insoluble debris was cleared by centrifugation at 12000 xg for 5 min at 4 °C. The sonicated chromatin (supernatant) was transferred to a new tube and stored at -80 °C.

For chromatin immunoprecipitation (ChIP), 25-100 µL Dynabeads Protein G beads (10003D or 10004D, Thermo Fisher Scientific) were washed twice with PBS-T (0.02% Tween-20 in 1x PBS). The desired amounts of antibodies (**Table S5**) were conjugated to the washed beads for 2-6 h at 4 °C with rotation. In parallel, frozen chromatin was thawed on ice, and the appropriate amount of chromatin (**Table S5**, determined by DNA content) was diluted with IP Dilution Buffer at a 1:4 ratio. Antibody-conjugated Dynabeads were washed twice with IP Dilution Buffer, resuspended in the diluted chromatin, and incubated overnight at 4 °C with rotation. The following day, the antibody-chromatin solution was cleared and the beads were washed four times with the following buffers: FA Lysis Buffer (50 mM HEPES-KOH [pH 7.5], 150 mM NaCl, 2 mM EDTA, 1% Triton X-100, 0.1% sodium deoxycholate, and 1x Complete Protease Inhibitor, EDTA-free); NaCl Buffer (50 mM HEPES-KOH [pH 7.5], 500 mM NaCl, 2 mM EDTA, 1% Triton X-100, and 0.1% sodium deoxycholate); LiCl Buffer (100 mM Tris-HCl [pH 8.0], 500 mM LiCl, 1% NP-40, and 1% sodium deoxycholate); and 10 mM Tris-HCl (pH 8.0). Chromatin was eluted with TES buffer (50 mM Tris-HCl [pH 8.0], 10 mM EDTA, and 1% SDS) for 45 min (15 min x 3) at 65 °C with shaking at 1000 rpm in a ThermoMixer F1.5 (EP5384000020, Eppendorf).

ChIP and Input samples were reverse crosslinked by adding NaCl to a final concentration of 200 mM and incubating for 2-8 h at 65 °C. Samples were treated with RNase A (50 ng/µL; EN0531, Thermo Fisher Scientific) for 0.5-1 h at 37 °C, followed by proteinase K treatment (0.2 mg/mL; 03115828001, Roche) for 2-4 h at 37 °C. DNA was purified by phenol-chloroform extraction followed by ethanol precipitation, and DNA concentration was measured using a Quantus fluorometer (E6150, Promega). Antibody and chromatin information are listed in **Table S5**.

### MNase assay

MNase-treated DNA samples were prepared following previously published protocols.^53,54^ Briefly, DE cells were dissociated and washed twice with Wash Buffer (20 mM HEPES-NaOH [pH 7.5], 150 mM NaCl, 0.5 M spermidine, and 1x Complete Protease Inhibitor, EDTA-free), then centrifuged at 600 xg for 3 min. The cell pellet was resuspended in Digitonin Buffer (0.02% digitonin in Wash Buffer). The cell suspension was then divided into aliquots of 200,000 cells per reaction and incubated with 2 U/mL (Low) or 80 U/mL (High) MNase (LS004798, Worthington) for exactly 5 min at 37 °C with shaking at 300 rpm in a ThermoMixer F1.5 (EP5384000020, Eppendorf). The MNase reaction was quenched by adding equal volumes of 2x Stop Buffer (340 mM NaCl, 20 mM EDTA, 4 mM EGTA, 0.02% digitonin, 0.05 mg/mL RNase A, and 0.05 mg/mL glycogen). For RNase treatment, samples were incubated with RNase A (EN0531, Thermo Fisher Scientific) at 37 °C for 30 min. Subsequently, samples were treated with 0.5% SDS and 0.2 mg/mL proteinase K for 2 h at 37 °C. Finally, DNA was purified by phenol-chloroform extraction followed by ethanol precipitation. The DNA concentration was measured using a NanoDrop spectrophotometer (13-400-518, Thermo Fisher Scientific). Mono- and sub-nucleosomal DNA fragments were selected and purified using MagBind Magnetic Beads (M1378-01, Omega). The purified DNA concentration was measured using a Quantus fluorometer (E6150, Promega).

### Library preparation and sequencing for ChIP-seq and MNase-seq

Multiplexed libraries of ChIP and MNase-treated samples were prepared from two replicates (corresponding to two independent CRISPRi clones) using the NEBNext Ultra II DNA Library Prep Kit for Illumina (E7645, NEB). We followed the manufacturer’s protocol with minor modifications: (1) after adaptor ligation, we performed library cleanup using AMPure XP Beads (A63880, Beckman Coulter) or MagBind Magnetic Beads (M1378-01, Omega) without size selection; (2) after PCR enrichment of the adaptor-ligated libraries, we performed two rounds of size selection using MagBind Magnetic Beads (M1378-01, Omega).

### Bulk RNA sequencing

One million cultured human cells were dissociated into single cells using Accutase and mixed with 100,000 *Drosophila* Schneider 2 (S2) cells for a 10% spike-in control. Cells were pelleted and lysed using Aurum Total RNA Lysis Solution (7326802, Bio-Rad). Total RNA was extracted using the Aurum Total RNA Mini Kit (7326820, Bio-Rad) and submitted to Novogene for library preparation and paired-end 150-bp sequencing on an Illumina NovaSeq 6000 platform.

### Bulk RNA-seq data analysis

Sequencing reads were aligned to the hg38 and dm6 combined genome using the STAR aligner.^78^ Only uniquely and concordantly aligned read pairs were used for downstream analysis. Gene-level read counts were quantified using featureCounts^82^ with the options ‘-O --fracOverlap 0.8’. For spike-in-controlled analysis, reads mapped to *Drosophila* genes were quantified using the same approach. Differential gene expression analysis was performed using the Wald test implemented in DESeq2.^83^ Spike-in normalization was incorporated by calculating size factors using the ‘estimateSizeFactors’ function in DESeq2 based on *Drosophila* gene read counts. Human genes with fold change (FC) ≥ 1.5 and false discovery rate (FDR) ≤ 0.05 were defined as differentially expressed genes for downstream analysis. Gene ontology (GO) analysis was performed using Enrichr,^84^ and the top significantly enriched GO terms were selected based on Benjamini-Hochberg-adjusted P values (derived from Fisher’s exact test) for presentation. Enriched GO terms were further summarized and visualized using the rrvgo R package.^96^ Bigwig files were generated on an RPM scale by pooling replicates per condition.

### ChIP-seq and MNase-seq data analysis

Sequencing reads were aligned to UCSC hg38 using the STAR aligner. Only uniquely aligned reads were retained for downstream analysis. Redundant reads were deduplicated per sample using the Picard tool.^97^ Biological replicates were pooled per condition and target to generate representative feature and bigwig files for presentation. Representative peaks of PRDM1 and pioneer transcription factors (pTF) were called against matching inputs by pooling replicates per condition. After peak calling, peaks with > 1 RPM (for pTFs) or > 0.7 RPM (for PRDM1) were retained for the downstream analysis. For MPRA design, PRDM1 and pTF peaks were called in a two-step process to leverage the statistical power of biological replicates. First, peaks were called against a matching input per replicate using Homer in a factor mode with default options. Peaks from replicates were pooled and merged into a candidate peak set. Anchoring on the candidate peaks, read counts were measured across ChIP and Input samples, which were subject to differential analysis using EdgeR. PRDM1 and PFs peaks were called with fold change ≥ 2, FDR ≤ 0.05, and > 0.7 RPM. Histone ChIP-seq data were also processed similarly, and bigwig files were generated on an RPM scale by pooling replicates per condition.

### ChromHMM data analysis

ChromHMM tracks from hPSC-derived endoderm cells were obtained from the Roadmap Epigenomics Mapping Consortium (https://egg2.wustl.edu/roadmap/web_portal, epigenome ID E011, hg38 lift-over) and simplified from an expanded 18-state into an 8-state model by collapsing related chromatin states with similar functional annotations as follows: TssA (TssA, TssAFlnk, TssFlnkU and TssFlnkD); GeneB (Tx and TxWk); EnhA (EnhA1 and EnhA2); EnhWk (EnhG1, EnhG2 and EnhWk); Het (ZNF/Rpts and Het); TssBiv (TssBiv); EnhPC (EnhBiv, RepPC and RepPCWk) and Quies (Quies).

### CellNet data training and query analysis

CellNet training and assessment of query samples was performed as previously described.^98^ Briefly, we mined NCBI GEO for bulk RNA-seq profiles of human embryonic stem cell derived samples from nine developmental trajectories: Human Pluripotent Stem Cells (hPSC), Anterior Primitive Streak (APS), Mid Primitive Streak (MPS), Definitive Endoderm (DE), Paraxial Mesoderm (PM), Lateral Plate Mesoderm (LPM), Neuroectoderm (NE) and Primordial Germ Cells (PGC). In total, more than 550 samples were collected across these cell types. FASTQ files were preprocessed into gene-by-sample expression matrices, quantile normalized and used as input to the gene regulatory network (GRN) reconstruction algorithm. GRN nodes corresponding to each reference cell type were then used as predictor variables to train a multi-class random forest (RF) classifier, following a top-scoring pair (TSP) gene-pair transformation that captures relative gene expression relationships. Classifier performance was evaluated using precision-recall (PR) curves, summarized as area under the PR curve (AUPR), using held-out subsets of the training data. Datasets used for CellNet training and validation are listed in **Table S6**.

To evaluate the impact of PRDM1 knockdown on endoderm lineage identity, bulk RNA-seq profiles from control and *PRDM1*-CRISPRi cell lines at the DE stage were processed using the gene filtering, normalization, and rank-transformation procedures implemented in CellNet. Then, query samples were assessed against the CellNet classifiers to generate identity scores for cell types/tissues originated from different developmental trajectories.

### lentiMPRA library design

The lentiMPRA library was generated using custom single-stranded DNA (ssDNA) oligonucleotide synthesis from Twist Bioscience and consisted of 15,336 oligos. Each oligo contained a 220 bp cis-regulatory sequence (CRS) from combinatorial mutations of TF motifs or scramble, flanked by two 15 bp adaptor sequences at the 5′ and 3′ ends, resulting in a total length of 250 bp. CRS were designed with the MPRA-ChIP designer pipeline (https://github.com/hwlim/MPRA-ChIP) as follows. Anchor regions were selected based on a subset of all PRDM1 ChIP-seq peaks at the DE stage, each centered and resized to 220 bp. PRDM1, EOMES, GATA, and SOX17 motifs (HOCOMOCO v11) were scanned with FIMO, retaining all hits at *P* ≤ 1e-4 or, when none was found, the single best hit at *P* ≤ 1e-3. Motif instances were intersected with the corresponding TF ChIP-seq co-binding table so that only motifs supported by ChIP-seq co-binding peaks were kept; anchors lacking a supported motif were discarded, and anchors bound by PRDM1 alone were randomly downsampled to 500, leaving a total of 2,012 anchors. For each anchor, we generated the wild-type (WT) sequence, combinatorial mutants (MUT) where each motif match was disrupted by shuffling nucleotides within the matched window, and a scramble (SCRAM) control where nucleotides from the WT sequences were randomly shuffled. Designed sequences were re-scanned with FIMO to confirm loss of targeted motif scores and preservation of non-targeted motifs. In total, the lentiMPRA library comprised 15,336 candidate CRSs, including 2,012 WT sequences; 2,012 SCRAM sequences; and 11,312 MUT sequences, in which one (n = 4,728), two (n = 4,287), three (n = 1,934), or four (n = 363) PRDM1 and/or pTF motifs were combinatorially mutated. The final designed oligo sequences are available in **Table S7.**

### lentiMPRA library cloning and sequence-barcode association

The lentiMPRA plasmid library was constructed as previously described,^45^ with minor modifications. Briefly, the array-synthesized oligonucleotide pool was first amplified by a low-cycle (5-cycle) PCR using primers (5BC-AG-f01 and 5BC-AG-r01_SV40; **Table S8**) designed to append an SV40 promoter and spacer sequence downstream of each candidate CRS. PCR products were purified using 1.8x MagBind Magnetic Beads (M1378-01, Omega) and subjected to a second amplification step consisting of 12 PCR cycles with primers (5BC-AG-f02 and 5BC-AG-r02; **Table S8**) that introduce a 15-nucleotide random barcode sequence. The barcoded amplicons were cloned into the SbfI/AgeI-digested pLS-SceI backbone (Addgene plasmid #137725) using NEBuilder HiFi DNA Assembly Mix (E2621S, NEB). The assembled library was electroporated into 10-beta electrocompetent E. coli (C3020, NEB) using a MicroPulser Electroporator (1652100, Bio-Rad), and transformants were grown overnight on carbenicillin-containing agar plates. Plasmid DNA was isolated by Midiprep (D4201, Zymo Research) from an estimated ∼1 million independent colonies, corresponding to an average representation of approximately 100 unique barcodes per CRS.

To map barcode–CRS associations, the CRS-SV40-barcode region was PCR-amplified from the pooled plasmid library using primers containing Illumina flow-cell adapters (P7-pLSmP-ass-gfp and P5-pLSmP-ass-i#; **Table S8**). The resulting amplicons were sequenced on a NextSeq X Plus platform using 150-bp paired-end reads with custom sequencing primers (R1: pLSmP-ass-seq-R1; index read: pLSmP-ass-seq-ind1; R2: pLSmP-ass-seq-R2_SV40; **Table S7**) to obtain ∼150 million total reads.

### Lentiviral infection and lentiMPRA barcode sequencing

Lentivirus for the MPRA was produced as previously described.^45^ Briefly, 10-12 million HEK293T cells were seeded in a single T225 flask. Two days later, plasmid DNA (10 µg plasmid library, 6.5 µg psPAX2, and 3.5 µg pMD2.G) was diluted in 800 µL Opti-MEM (31985062, Thermo Fisher Scientific) to generate premix A. In parallel, 60 µL EndoFectin (EF001, GeneCopoeia) was diluted in 800 µL Opti-MEM to generate premix B. Premixes A and B were combined, incubated at room temperature for 15 min, and then added dropwise to the cells. Cells were incubated for 8-14 h, after which the medium was replaced with 30 mL per flask of DMEM supplemented with 5% heat-inactivated FBS and 1x ViralBoost reagent (VB100, Alstem; 60 µL per 30 mL medium). Cells were incubated for an additional 48 h, after which lentiviral supernatant was collected, filtered through a 0.45-µm PES filter system (165-0045, Thermo Scientific), and concentrated using Lenti-X Concentrator (631232, Takara Bio).

Titration of the lentiMPRA library was conducted on 72.3 hPSC lines as previously described,^45^ with minor modifications. Briefly, hPSC cells were plated at 1 × 10^6^ cells per well in 6-well plates and incubated for 24 h. The following day, cells were transduced with serial volumes of lentivirus (0, 4, 8, 16, 32, or 64 µL) in medium supplemented with 8 µg/mL polybrene and spin-infected by centrifugation at 930 x g for 90 min at 32 °C. Transduction medium was replaced with fresh medium after 4 – 8 h, and cells were cultured for 3 days prior to genomic DNA extraction. Genomic DNA was extracted using the Wizard SV genomic DNA purification kit (A2361, Promega). The multiplicity of infection (MOI) was measured as the relative abundance of viral DNA (WPRE region, WPRE.F and WPRE.R) over that of genomic DNA [intronic region of LIPC gene, LP34.F and LP34.R (**Table S8**)] by qPCR using the PowerUp™ SYBR™ Green Master Mix (A25778, Thermo Fisher Scientific), according to the manufacturer’s protocol.

Lentiviral infection, DNA/RNA extraction, and barcode sequencing were all performed as previously described,^45^ with minor modifications. Briefly, ∼14 million cells (two 6-well plates) per replicate were infected with the lentiviral library at a MOI of 8 in medium supplemented with 8 µg/mL polybrene (TR-1003-G, Sigma-Aldrich). Three independent biological replicates were generated. One day after transduction, cells were differentiated to anterior primitive streak (APS), followed by differentiation to definitive endoderm (DE) the next day. On day 3, cells were harvested, and DNA and RNA were purified using TRIzol™ Reagent (15596026, Thermo Fisher Scientific). RNA was treated with TURBO DNA-free™ Kit (AM1907, Thermo Fisher Scientific) to remove contaminating DNA, and reverse-transcribed with SuperScript II (18064022, Thermo Fisher Scientific) using a barcode-specific primer containing a unique molecular identifier (UMI) (P7-pLSmp-assUMI-gfp, **Table S8**). Barcode DNA or cDNA from each replicate was amplified with a 3-cycle PCR using specific primers (P7-pLSmp-assUMI-gfp and P5-pLSmP-5bc-i#, **Table S8**) to add the sample index and UMI. PCR products were then subjected to a second amplification step consisting of 16 cycles using P5 and P7 primers (**Table S8**). Amplified fragments were purified and sequenced on an Illumina NovaSeq X Plus platform using 15-bp paired-end reads and 10-cycle dual-index reads with custom sequencing primers (R1, pLSmP-ass-seq-ind1; R2 (index read1 for UMI), pLSmP-UMI-seq; R3, pLSmP-bc-seq; R4 (index read2 for sample index), pLSmP-5bc-seqR2, **Table S8**).

### lentiMPRA data analysis

Association between CRS and barcode was done using the MPRAflow package. Given the homology between CRS, we revised the default association process in MPRAflow to consider only perfect matches. The CRS–barcode association results were provided to the count process of the MPRAflow to construct DNA and RNA readout counts per associated barcode.

Count matrices were reformatted for use as input to MPRAnalyze. Specifically, barcode-level DNA and RNA count matrices and their associated column annotations were generated with the MPRAflow and imported into R, with missing entries set to zero. Because non-zero counts were concentrated in the first barcode positions of each replicate, the data were downsampled to the first 50 barcodes per replicate across all four replicates (200 barcode–replicate units in total), and the count matrices and annotation tables were truncated accordingly. Sparsity before and after downsampling was inspected by heatmaps of the DNA and RNA. CRS metadata were then restricted to CRSs present in the count data. Each CRS was defined by its anchor region and the transcription factor motif that had been mutated within it, with wild-type and scrambled sequences treated as additional motif categories. To enable comparisons among mutants sharing an anchor, the barcode-level matrices were restructured into matrices in which rows corresponded to anchors and columns to the full combination of mutation, replicate, and barcode; each column was annotated with its mutation identity and classified as wild-type, scrambled, or test. Counts for each CRS were placed in the row of its anchor, the column block of its mutation, and column assignments were verified against the mutation labels. DNA and RNA matrices were processed identically and, together with their annotation tables, were used as input for activity quantification and differential activity analysis with MPRAnalyze.

For every model fit in MPRAnalyze, DNA and RNA library depth factors were estimated jointly with estimateDepthFactors() using replicate as the library factor and the upper-quartile depth estimator. The whole workflow was run three times, identically except for the control (reference) level used as the model baseline: WT, SCRAM, or PRDM1-MUT. For quantification, analyzeQuantification() was used, fitting the nested DNA and RNA gamma-Poisson models with dnaDesign = ∼ Mutation and rnaDesign = ∼ Mutation. Barcode was deliberately excluded from the quantification design, since per-barcode activity estimates are not comparable across loci. Activity was extracted as the transcription-rate parameter α with getAlpha(by.factor = “Mutation”). Differential activity was assessed pairwise, comparing each of the 32 non-reference mutations against the control in a separate two-level model. This pairwise formulation permits an allelic design: because each mutation carries its own barcode set, barcode is nested within mutation, and DNA copy number was modeled per oligo using barcode_allelic = interaction(barcode, Mutation). For each pair, counts and annotations were subset to the two conditions, the control was set as the reference level, and analyzeComparative() was fitted in classic mode with dnaDesign = ∼ replicate + barcode_allelic, rnaDesign = ∼ Mutation, and fit.se = TRUE. Coefficients were tested with testCoefficient(factor = “Mutation”), and the natural-log fold changes were converted to log2. The resulting per-locus log2 fold change, test statistic, *P*-value, and Benjamini–Hochberg FDR were collected across all 32 comparisons. A locus was called differentially active for a given mutation when FDR ≤ 0.05.

## QUANTIFICATION AND STATISTICAL ANALYSIS

### Mixed-effects modeling of lentiMPRA activity

Analyses were conducted in R using the lme4, lmerTest and emmeans packages. Differences in MPRA activity were assessed using linear mixed-effects models with Anchor included as a random intercept to account for repeated measurements across constructs derived from the same CRS. A linear trend model was fit as logFC ∼ motif_count * PRDM1_group + (1 | Anchor), and a categorical model was fit as logFC ∼ factor(motif_count) * PRDM1_group + (1 | Anchor) to estimate activity at each motif class. Estimated marginal means and pairwise contrasts were computed using emmeans, with Kenward–Roger degrees of freedom for inference.

---

**Figure S1.**
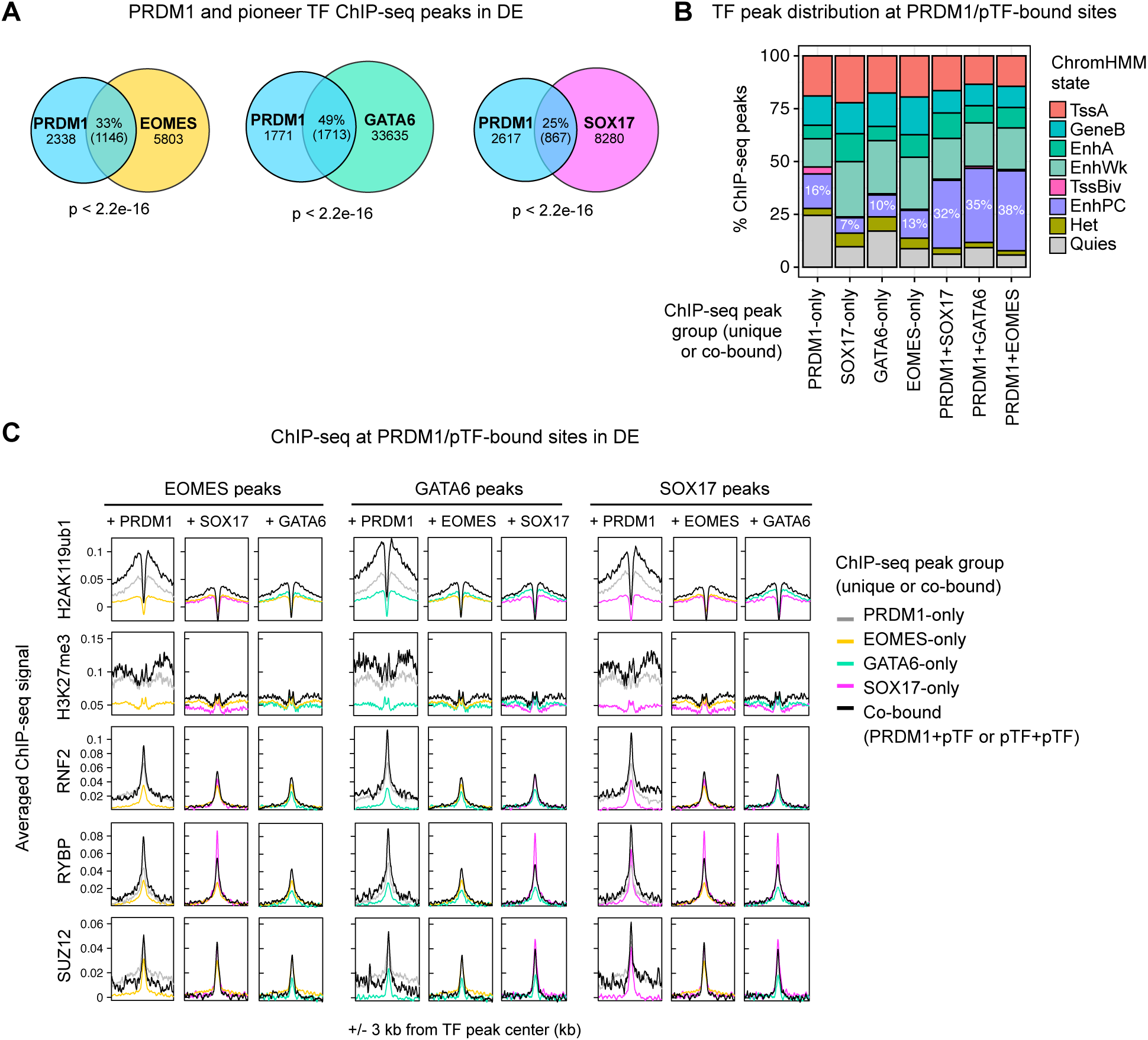
PRDM1 cooperates with stage- and lineage-specific pioneer TFs to establish Polycomb-associated enhancers, related to Figure 1. **(A)** Venn diagrams showing pairwise intersections between PRDM1 and GATA6, EOMES, and SOX17 ChIP-seq peaks in DE cells. Numbers indicate unique and shared peak counts. *P-*values were calculated using Fisher’s exact test to assess whether the observed overlaps exceeded those expected by chance. **(B)** Bar plots showing the percentage of ChIP-seq peaks assigned to each ChromHMM state, relative to the total number of peaks in each chromatin state. **(C)** Averaged ChIP-seq signal profiles centered on unique- or co-bound TF peaks showing enrichment of PRC1 and PRC2 subunits (RING1B, RYBP, and SUZ12) and their associated histone modifications (H2AK119ub1 and H3K27me3) in DE cells.

**Figure S2.**
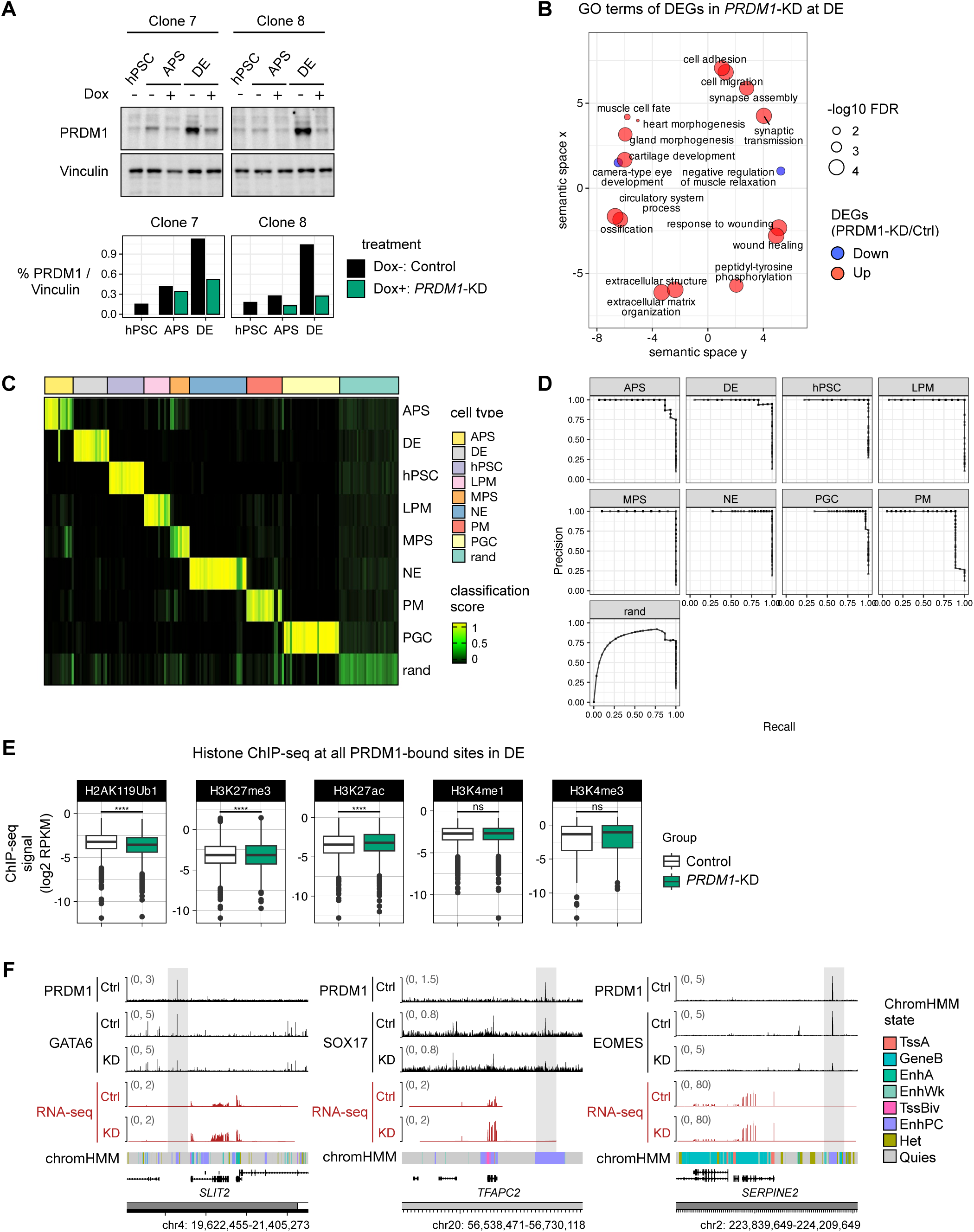
PRDM1 safeguards endoderm identity by Polycomb-mediated repression of alternative-lineage genes, related to Figure 2. **(A)** Assessment of *PRDM1*-CRISPRi knockdown (KD) efficiency by Western blotting during differentiation from hPSC to DE stages (*n* = 2 biological replicates). **(B)** Gene Ontology (GO) terms for differentially expressed genes (DEGs) in *PRDM1*-KD cells at the DE stage. **(C)** Classification heatmap of the RNA-seq dataset. CellNet binary classifiers were trained for each cell type using cell type-specific gene regulatory network (GRN) genes. Each row represents a cell type classifier, and each column represents a validation sample. Higher classification scores indicate greater similarity to the corresponding reference cell type. APS, anterior primitive streak; MPS, mid primitive streak; PGC, primordial germ cell; hPSC, human pluripotent stem cell; DE, definitive endoderm; LPM, lateral plate mesoderm; PM, paraxial mesoderm; NE, neuroectoderm; rand, random. **(D)** Precision-recall curves evaluating the performance of CellNet classifiers on the validation dataset. Precision measures the fraction of predicted positive samples that are correctly classified, whereas recall measures the fraction of true positive samples that are correctly identified. Higher precision and recall indicate better classifier performance. **(E)** Box plots of ChIP-seq signal at +/- 3 kb from PRDM1 peaks for active (H3K27ac), active/primed enhancer-associated (H3K4me1), active/primed promoter-associated (H3K4me3), PRC2-mediated (H3K27me3), and PRC1-mediated (H2AK119ub1) marks in control and *PRDM1*-KD DE cells. Statistical significance was assessed using the Wilcoxon rank-sum test to compare control and *PRDM1*-KD samples. Significance thresholds: *P* ≤ 0.05 (*), *P* ≤ 0.01 (**), *P* ≤ 0.001 (***); ns, not significant. **(F)** Genome browser tracks showing ChromHMM state annotations, TF ChIP-seq signal, and RNA-seq coverage in control and *PRDM1*-KD cells at representative loci in DE cells.

**Figure S3.**
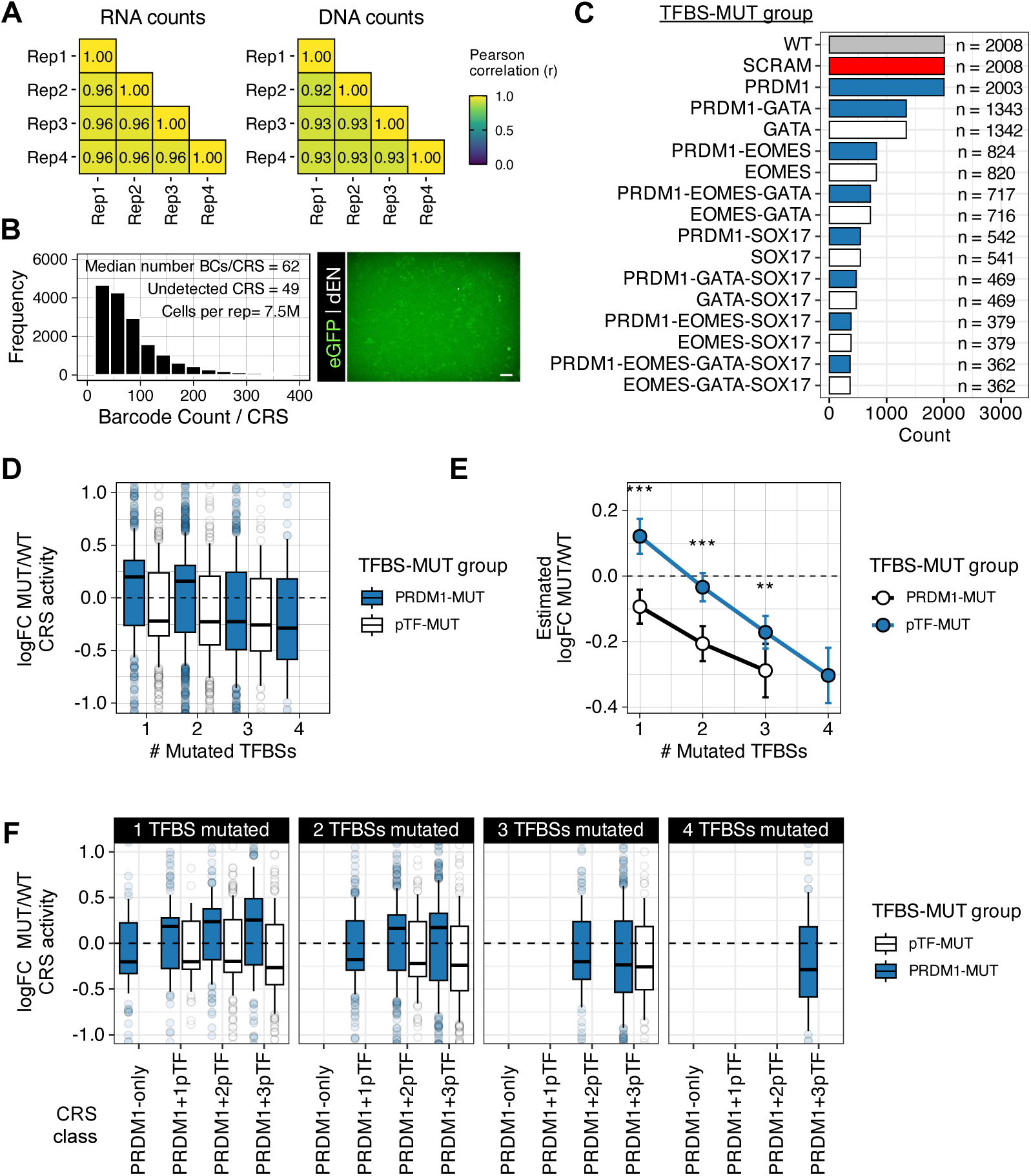
The number of pioneer TF-binding sites determines the strength of PRDM1-mediated enhancer repression, related to Figure 3. **(A)** Pairwise Pearson correlation matrices of log2-normalized RNA and DNA counts across biological replicates. Correlation coefficients (*R*) were calculated separately for RNA and DNA barcode counts to assess the reproducibility of library representation and transcriptional output across biological replicates. **(B)** Frequency distribution of DNA barcode counts per CRS, showing the number of unique barcodes associated with each CRS (left), and a representative image of DE cells transduced with the lentiMPRA library, showing EGFP reporter expression (right). The scale bar is 100 µm. **(C)** Bar plot showing the number of detected CRSs across WT (grey), SCRAM (red), PRDM1-PERT (blue), and pTF-PERT (white) groups following transduction of the lentiMPRA library. Names on the y-axis correspond to the TFBSs mutated in that specific CRS group. **(D)** Box plots showing fold changes in CRS activity between TFBS-MUT constructs and corresponding WT sequences across mutation groups, stratified by the number of mutated TFBSs. **(E)** Estimated marginal means for fold changes in CRS activity between TFBS-MUT constructs and corresponding WT sequences across mutation groups, PRDM1-MUT and pTF-MUT. Error bars indicate 95% confidence intervals. Asterisks denote pairwise comparisons between PRDM1-MUT and pTF-MUT constructs within each motif class: *P* ≤ 0.05 (\**), P* ≤ 0.01 (**), *P* ≤ 0.001 (***); ns, not significant. **(F)** Box plots showing fold changes in CRS activity between TFBS-MUT constructs and corresponding WT sequences, across CRS classes, stratified by the number of TFBSs.

**Figure S4.**
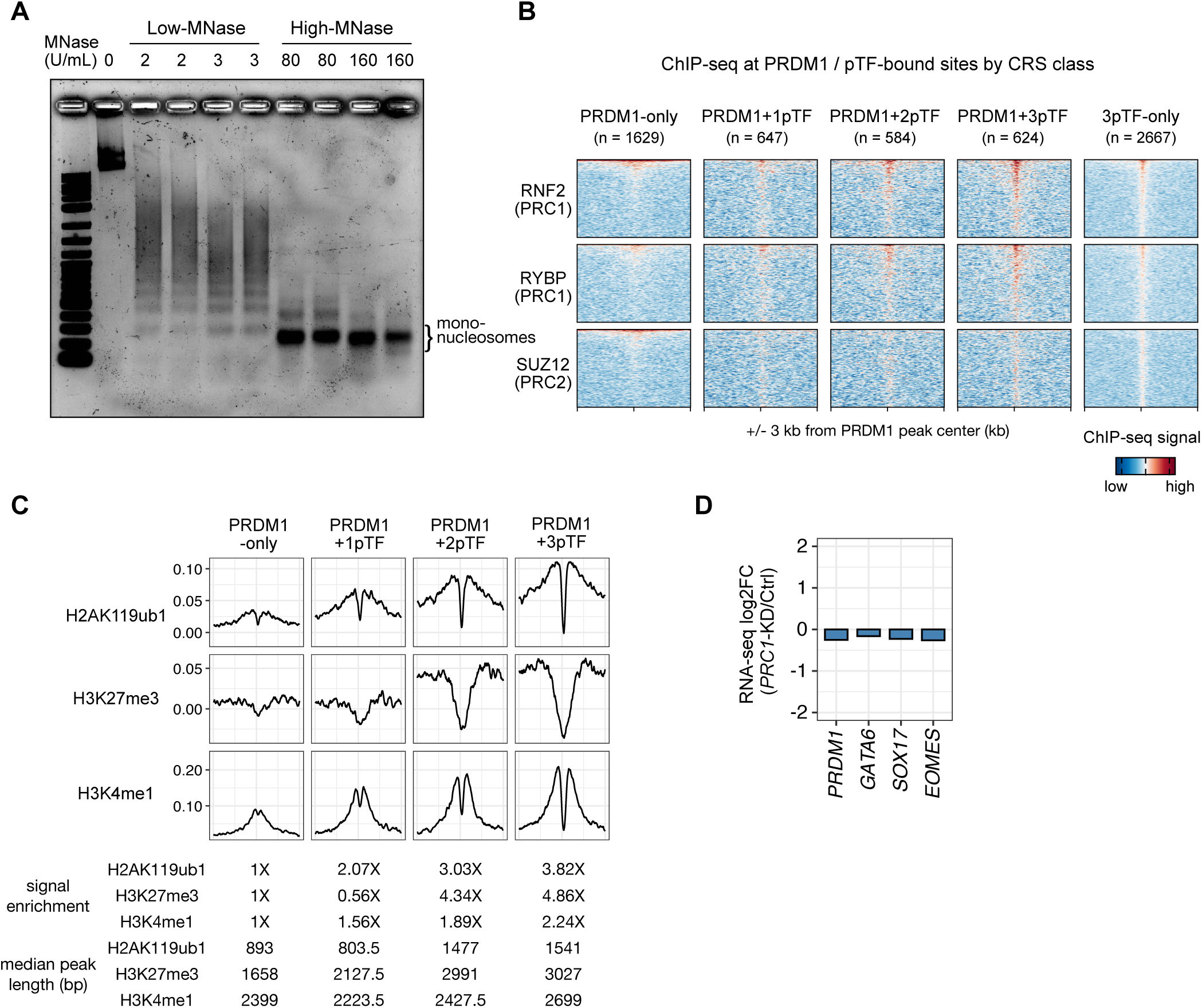
Collaborative binding by diverse pioneer TFs and PRDM1 establishes focal chromatin accessibility and promotes hyper-bivalent enhancer formation, related to Figure 4. **(A)** Agarose gel showing DNA fragments following Low- and High-MNase digestion. Increased MNase concentration resulted in greater digestion and enrichment of mono-nucleosomal fragments. **(B)** Heatmaps of ChIP-seq signal for PRC subunits centered on PRDM1 peaks or 3pTF-only peaks across CRS classes. **(C)** Averaged ChIP-seq signal profiles and quantifications for bivalent modifications (H2AK119ub1, H3K27me3, and H3K4me1) at enhancers centered on PRDM1 peaks in DE cells. **(D)** Expression changes for *PRDM1* and *pioneer TF* genes in *PRC1*-KD DE cells compared to control cells.

**Table S1.** RNA-seq differential expression analysis of hPSCs at the hPSC, ME, DE, PFG, and LBP stages

**Table S2.** RNA-seq differential expression analysis of *PRDM1*-CRISPRi at the DE stage

**Table S3.** Antibody information for Western blot assays.

| Antigen | Supplier | Cat. No. | Lot No. | Host | Dilution factor |
| --- | --- | --- | --- | --- | --- |
| PRDM1 | Cell Signaling Technology | 9115S | 6, 7 | Rabbit | 1:1000 |
| RNF2/ RING1B | Bethyl | A302-869A | 1 | Rabbit | 1:1000 |
| RYBP | Millipore | AB3637 | 3827461 | Rabbit | 1:1000 |
| SUZ12 | Cell Signaling Technology | 3737S | 8 | Rabbit | 1:1000 |
| VINCULIN | Sigma-Aldrich | V-9131 | 118M4777V | Mouse | 1:5000 |
| TrueBlot Anti-Goat IgG_HRP | Rockland | 18-8814-31 | 46356 | Mouse | 1:1000 |
| TrueBlot Anti-Rabbit IgG_HRP | Rockland | 18-8816-33 | 43708 | Mouse | 1:1000 |
| StarBright™ Blue 700 Goat Anti-Rabbit IgG | Bio-Rad | 12004162 | 64297092 | Goat | 1:5000 |
| StarBright™ Blue 520 Goat Anti-Mouse IgG | Bio-Rad | 12005867 | 64264264 | Goat | 1:5000 |

**Table S4.** Antibody information for co-IP experiments.

| Antigen | Supplier | Cat. No. | Lot No. | Host | Antibody amount |
| --- | --- | --- | --- | --- | --- |
| PRDM1 | Cell Signaling Technology | 9115S | 6, 7 | Rabbit | 0.1 ug |
| GATA6 | Cell Signaling Technology | 5851S | 5, 6 | Rabbit | 5.0 ug |
| SOX17 | R&D Systems | AF1924 | KGA1022031 | Goat | 5.0 ug |
| EOMES | Abcam | Ab23345 | GR3442174-1 | Rabbit | 5.0 ug |
| RING1B/RNF2 | Bethyl | A302-869A | 1 | Rabbit | 4.0 ug |
| RYBP | Millipore | AB3637 | 3827461 | Rabbit | 5.0 ug |
| SUZ12 | Cell Signaling Technology | 3737S | 8 | Rabbit | 0.5 ug |
| Normal Goat IgG | R&D Systems | AB-108-C | ES4521041 | Goat | 4.0 ug |
| Normal Rabbit IgG | Millipore | 12-370 | 3059594 | Rabbit | 4.0 ug |

**Table S5.** Antibody and chromatin information for ChIP experiments.

| Antigen | Supplier | Cat. No. | Lot No. | Host | Antibody amount | Chromatin (DNA) amount |
| --- | --- | --- | --- | --- | --- | --- |
| PRDM1 | Cell Signaling Technology | 9115S | 6, 7 | Rabbit | 0.1 ug | 75.0 ug |
| GATA6 | Cell Signaling Technology | 5851S | 5, 6 | Rabbit | 5.0 ug | 50.0 ug |
| SOX17 | R&D Systems | AF1924 | KGA1022031 | Goat | 5.0 ug | 50.0 ug |
| EOMES | Abcam | Ab23345 | GR3442174-1 | Rabbit | 5.0 ug | 50.0 ug |
| RING1B/RNF2 | Bethyl | A302-869A | 1 | Rabbit | 3.0 ug | 37.5 ug |
| RYBP | Millipore | AB3637 | 3827461 | Rabbit | 4.0 ug | 37.5 ug |
| SUZ12 | Cell Signaling Technology | 3737S | 8 | Rabbit | 0.4 ug | 37.5 ug |
| H2AK119ub1 | Cell Signaling Technology | 8240S | 8, 9 | Rabbit | 2.5 ug | 18.75 ug |
| H3K27me3 | Cell Signaling Technology | 9733S | 14 | Rabbit | 0.5 ug | 25.0 ug |
| H3K27ac | Abcam | ab4729 | GR1033973-1 | Rabbit | 2.5 ug | 18.75 ug |
| H3K9me3 | Abcam | ab8898 | GR106377-01 | Rabbit | 2.5 ug | 3.0 ug |
| H3K9me2 | Abcam | ab1220 | GR3247768-1 | Mouse | 5.0 ug | 10.0 ug |
| H3K4me1 | Abcam | ab8895 | GR3206758-1 | Rabbit | 2.0 ug | 18.75 ug |
| H3K4me3 | Millipore | 07-473 | 3191394 | Rabbit | 2.0 ug | 18.75 ug |

**Table S6.** CellNet training and validation datasets

**Table S7.** lentiMPRA anchor regions, motif masking, and CRS-barcode associations

**Table S8.**
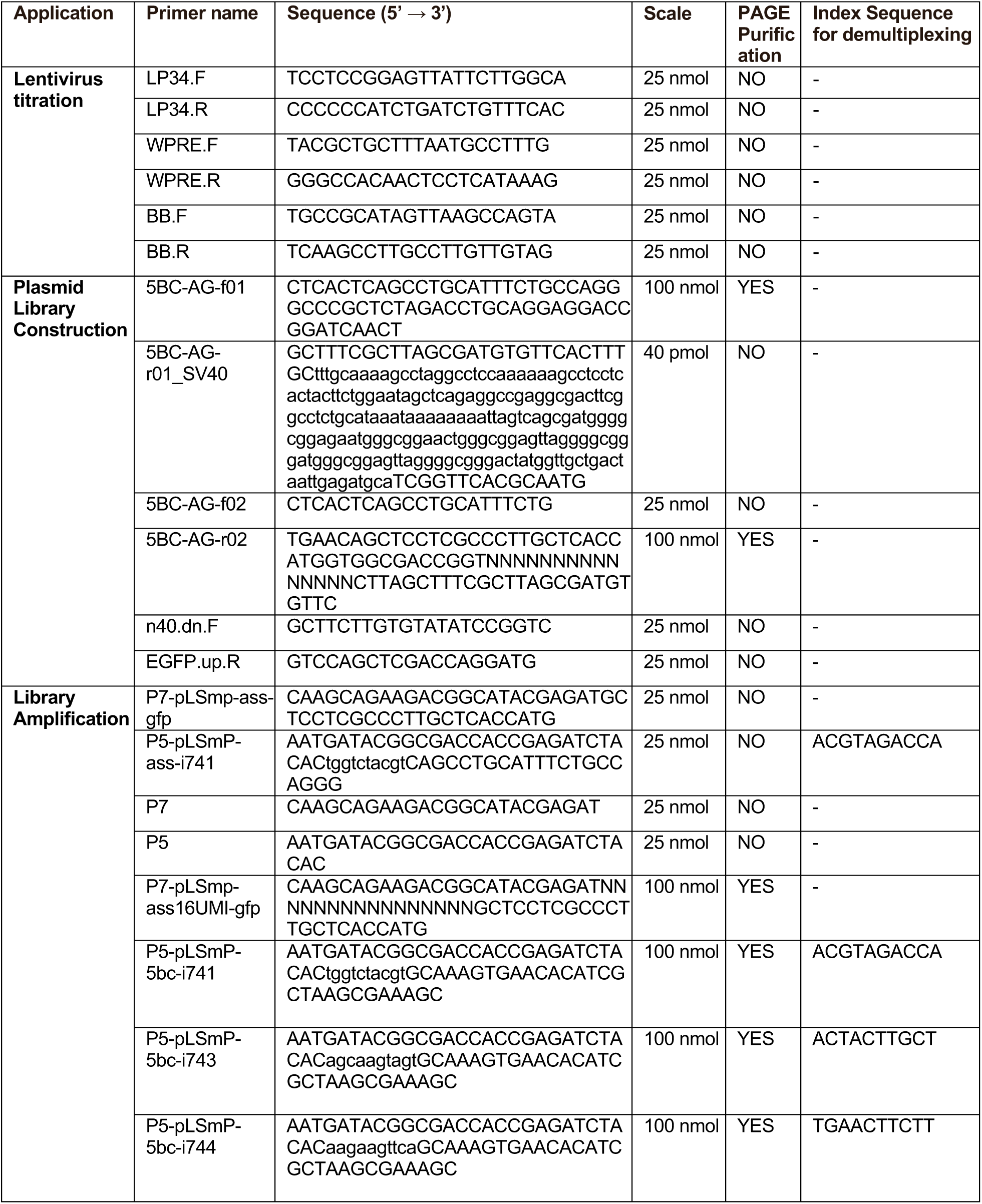
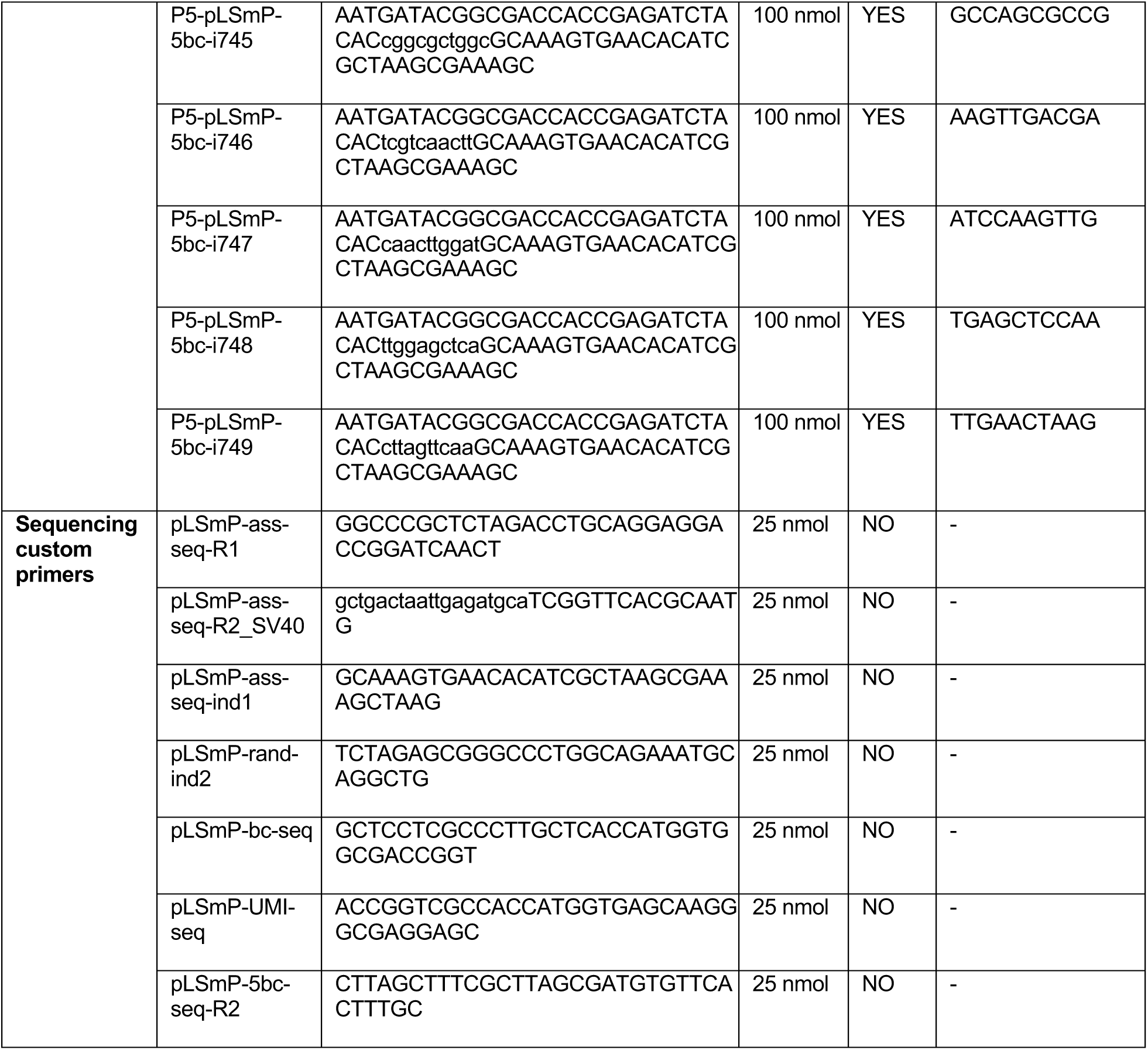
Primer sequences used for lentiMPRA.

